# Isoform switch of CD47 provokes macrophage-mediated pyroptosis in ovarian cancer

**DOI:** 10.1101/2025.04.17.649282

**Authors:** Zixiang Wang, Lei Yang, Siyuan Yang, Gaoyuan Li, Meining Xu, Beihua Kong, Changshun Shao, Zhaojian Liu

**Author notes:** Dr. **Zhaojian Liu**; Dr. **Changshun Shao**; Dr. **Beihua Kong**. These authors contributed equally to this work (Zixiang Wang, Lei Yang).

## Abstract

Cancer cells expressing CD47 escape macrophage phagocytosis by binding to the SIRPα ligand expressed on macrophages. CD47 targeted therapy offers a promising approach to cancer treatment. Here we report that two major isoforms of CD47 differ in ovarian cancer and normal tissues. The truncated isoform lacking exon 9 and exon 10 (CD47-S) was exclusively expressed in normal tissues, whereas the full-length isoform (CD47-L) was predominantly expressed in ovarian cancer tissues. Interestingly, CD47-S was unable to locate at the cell surface to bind with SIRPα as CD47-L did, and thereby inactivated “don’t-eat-me” signal. Mechanistic investigations revealed that splicing factor HNRNPA1 promoted splicing switch from *CD47-L* to *CD47-S* through exon 9 and exon 10 skipping. We further developed antisense oligonucleotides (ASOs) that effectively switched *CD47-L* to *CD47-S*. Importantly, ASOs treatment evoked macrophage-mediated antitumor immune response, thereby triggered pyroptosis of ovarian cancer cells. Moreover, CD47-targeting ASOs significantly reduced tumor growth in patient-derived xenograft. Together, ASO-mediated isoform switch of CD47 has emerged as a promising strategy to improve immune responses against tumors.

## Introduction

Ovarian cancer is one of the most lethal gynecologic malignancies, characterized by a poor prognosis and high mortality rates due to its typically late diagnosis and frequent recurrence.^1^ Despite the clinical success of immune checkpoint therapies (ICT) for the treatment of various human malignancies, the overall response rate remains relatively low in ovarian cancer.^2^ The overall objective responses of PD-1/PD-L1 inhibitors for ovarian cancer patients is less than 10%.^3^ There is no significant difference in overall survival between monotherapy of anti-PD-(L)1 and combination therapy of anti-CTLA4 and anti-PD-(L)1.^4^ There is an urgent need for developing effective immunotherapy strategies in ovarian cancer.

Accumulating evidence suggests that ovarian cancer has a highly immunosuppressive tumor microenvironment (TME) that mediated resistance to ICT. TME is associated with the recruitment of immunosuppressive myeloid-derived suppressor cells (MDSCs), regulatory T cells (Tregs) and tumor association macrophages (TAMs).^5^ In the ovarian cancer microenvironment, TAMs are tended to exhibit the M2-like phenotype and have been shown to participate in the pathogenesis and progression of ovarian cancer.^6^ TAMs from ovarian ascites have been shown to suppress the function of NK cell and T cells.^7^ PD-1 expressing TAMs correlates negatively with phagocytic potency against tumor cells.^8^ Ovarian cancer cell-secreted interleukin-4 promotes resistance to anti-PD-1 through its effects on macrophages.^3^ Targeting tumor associated macrophages is a promising immunotherapeutic strategy for ovarian cancer.

CD47 is a transmembrane glycoprotein that functions as a “don’t eat me" signal when it binds SIRPα on macrophages, thereby protects cancer cells from macrophage-mediated phagocytosis.^9^ CD47 is overexpressed in various types of human malignancies, its high level is associated with poor prognosis and reduced therapeutic responses.^10^ Antibodies blocking CD47 have demonstrated potent anti-tumor activity of hematologic malignancies and solid tumors in preclinical models and clinical studies.^11^ Nevertheless, hematotoxicity has emerged as the most common adverse effect that limit their clinical application. In this study, we identified two distinct isoforms of CD47 selectively expressed in ovarian cancer and normal tissues. CD47-L, but not CD47-S is preferentially expressed in ovarian cancer cells associated with immunosuppressive tumor microenvironment. We further demonstrated that CD47-S was unable to locate at the cell surface to bind with SIRPα as CD47-L did. Importantly, we developed a splice-switch antisense oligonucleotide that successfully switch CD47-L to CD47-S, thereby evoked macrophage-mediated pyroptosis of ovarian cancer cells. Our findings suggest that targeting isoform switch of CD47 appears as a promising strategy to improve immune responses against tumors.

## Result

### CD47 isoforms differ in ovarian cancer and normal tissues

To identify immune checkpoint genes that aberrantly spliced in ovarian cancer, we performed PacBio long read RNA sequencing in 8 pairs of serous ovarian cancer and normal tissues, combined with RNA-seq data from TCGA-OV and normal ovary and fallopian tube epithelium, a total of 2761 ovarian cancer specific isoform switch genes with functional consequences were identified **(Fig. 1A**). We next integrated the 2761 genes with 32 dysregulated immune checkpoint ligands^12^ and obtained 7 genes, including CEACAM1, TNFSF14, CD47, BTN2A1, HLA-A, HLA-DMA, HLA-DMB **(Fig. 1B and C)**. CD47 was identified as one of the most differentially expressed and aberrantly spliced immune checkpoint genes.

**Figure 1.**
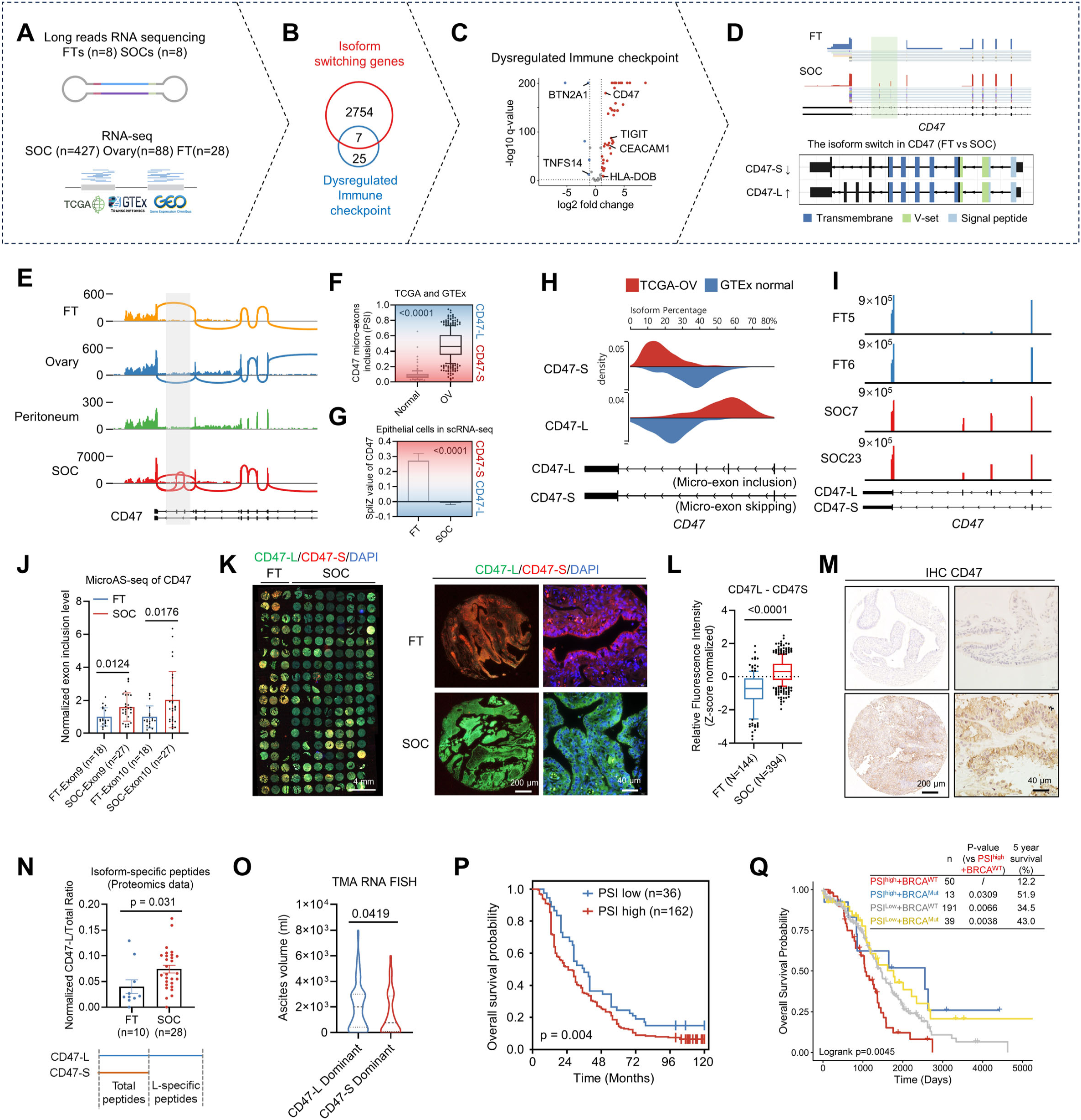
CD47 isoforms differ in ovarian cancer and normal tissues. **(A)** Diagram depicting the workflow for identifying immune checkpoints undergoing isoform switching in ovarian cancer. Long reads and short reads transcriptome data were used to analyze isoform switches. **(B)** Venn diagram indicates 7 dysregulated immune checkpoint ligands were detected isoform switch comparing SOCs with FTs. **(C)** Volcano plot of differential expression of immune checkpoints in TCGA-OV and GTEx-ovary. **(D)** Representative genomic visualization and schematic diagram of gene structure of CD47 based on long reads sequencing of the FT and SOC sample. **(E)** Representative genomic visualization of CD47 in FT, ovary, peritoneum, and SOC tissues. Junction between exons were shown as arc. **(F)** PSI index of CD47 exon 9 and 10 skipping in TCGA-OV and GTEx-ovary. **(G)** SpliZ value of epithelial cells in scRNA-seq data from FT and SOCs. **(H)** Isoform percentage of CD47-L and CD47-S in TCGA-OV and GTEx ovary database. **(I and J)** Genomic visualization **(I)** and quantification **(J)** of MicroAS-seq targeting CD47 micro-exon 9 and 10 skipping regions of FT and SOC samples. **(K)** Representative images of RNA FISH of CD47 isoforms detected by immunohistochemistry in TMA containing 144 FT and 394 SOC spots. **(L)** Normalized relative fluorescence intensity of CD47-L and CD47-S of samples in TMA. **(M)** Representative images of immunohistochemistry on the same microarray using anti-CD47 antibody. **(N)** Proportion of the CD47-L isoform in FTs (n=10) and SOCs (n=28) calculated based on peptide abundance from quantitative proteomics. **(O)** The ascites volume of patients with ovarian cancer in CD47-L and CD47-S dominant groups. **(P)** Prognostic analysis of the correlation between CD47 isoforms and overall survival of Qilu cohort. 198 patients were divided into PSI low (n=36) and high (n=162) group. **(Q)** Kaplan-Meier survival curves showing overall survival probability stratified by CD47 PSI expression levels (high or low) and BRCA mutation status (wild-type or mutant). The p value was obtained by log-rank test.

Our analysis on long read RNA sequencing data revealed that two micro-exons (exon 9 and exon 10) of CD47 were exclusively expressed in cancerous tissues **(Fig. 1D)**. Meanwhile, the inclusion level of exon 9 and exon 10 was significantly increased in ovarian cancer compared with normal tissues, especially in epithelial cells **(Fig. 1E-G).** Consequently, the short isoform lacking exon 9 and exon 10 (CD47-S) was predominantly expressed in normal tissues, whereas the long isoform with exon 9 and exon 10 inclusion (CD47-L) was exclusively expressed in ovarian cancer tissues **(Fig. 1H)**. Alternative micro exons (less than 51 nucleotides)^13^ are too short to be detected by semiquantitive-PCR, we developed microAS-seq, a high-throughput sequencing demultiplexing method using targeted amplification primers to quantify specific individual exon inclusion. Using this method, we found exon 9 and exon 10 inclusion level was higher in ovarian cancer compared with normal tissues (**Fig. 1I-J**). The conclusion was also verified with classic isoform specific qPCR. We next performed RNA FISH on a tissue microarray comprised of 144 normal fallopian tube (FT) and 394 serous ovarian cancer (SOC) spots using isoform specific probes to detect CD47-S (red) and CD47-L (green) respectively. As a result, the fluorescence intensity of CD47-L in SOC tissues was higher relative to FT tissues (**Fig. 1K-L**). By analyzing pan-cancer data from TCGA and GTEx, we observed inclusion of exon 9 and exon 10 was correlated with overexpression of overall level of CD47, especially in ovarian cancer. Immunohistochemistry on the same microarray using anti-CD47 antibody indicated that CD47 protein expression was correlated with CD47-L/S ratio, which was further validated in the 12 ovarian cancer cell lines from the Cancer Cell Line Encyclopedia (CCLE). In consistent, CD47 protein and RNA expression was significantly higher in SOC tissues (**Fig. 1M**). Furthermore, we conducted quantitative proteomics in FTs (n=10) and SOCs (n=28) tissues and found CD47-L specific peptides ratio was significantly upregulated in SOCs compared to FTs (**Fig. 1N**). Moreover, pan-cancer RNA sequencing data analysis showed that CD47 exon 9 and exon 10 inclusion level was higher in various cancers compared to normal tissues.

To assess the clinical relevance of CD47 isoform switch in ovarian cancer, we analyzed CD47-L expression on RNA FISH data in the TMA with 394 serous ovarian cancer tissues. Intriguingly, CD47-L was significantly correlated with poor prognosis, higher ascites volume and p53 positive ratio, and lower blood ferritin concentration in Qilu cohort **(Fig. 1O-P).** More importantly, we divided TCGA-OV cohort based on percent spliced in (PSI) level of CD47 exon 9 and exon 10 and BRCA1/2 mutation status. The CD47 PSI high and BRCA wild-type group showed worst prognosis with a 12.2% 5-year survival **(Fig. 1Q).** Collectively, these results indicate that two CD47 isoforms differ in ovarian cancer and normal tissues, CD47-L is the abundant isoform in cancerous tissues and its high level is associated with worse prognosis in ovarian cancer patients.

### CD47-L, but not CD47-S is preferentially expressed in ovarian cancer cells associated with immunosuppressive tumor microenvironment

We next sought to characterize the expression pattern of CD47-L and CD47-S in TME of ovarian cancer by analyzing scRNA-seq and spatial transcriptome data **(Fig. 2A).** We first integrated 5 public scRNA-seq datasets of 20 serous ovarian cancer (SOC) tumors and identified seven major cell types including NK/T cells, fibroblasts, myeloid cells, endothelial cells, plasma cells, B cells and epithelial cells. Interestingly, both CD47-L and CD47-S were observed in epithelial subtypes (**Fig. 2B-C).** We next conducted cell-cell communication analysis and found that CD47-S^high^ epithelial cells interacted more closely with NK/T cells and myeloid cells compared with CD47-L^high^ epithelial cells, marked by CD86, ICAM, ITGB2 and CD40 **(Fig. 2D-E)**. Further functional enrichment analysis suggested that CD47-S^high^ epithelial cells showed a robust enrichment of molecules involved in apoptosis and IFN-alpha response, whereas CD47-L^high^ epithelial cells showed a close correlation with hypoxia and TGF-beta signaling (**Fig. 2F)**. In parallel, we subdivided these 20 samples into CD47-S dominant samples and CD47-L dominant samples based on SpliZ values. Cell-Chat analysis revealed that CD47-S dominant samples had stronger communication strength with immune-active tumor microenvironment compared to CD47-L dominant samples (**Fig. 2G-H)**. Consistently, the RNA expression level of macrophage pro-inflammatory and T cell activation genes was significantly upregulated in CD47-S dominant samples compared to CD47-L dominant samples **(Fig. 2I-J)**. Furthermore, subpopulation analysis of myeloid cells revealed a higher proportion of pro-inflammatory subsets Macro_NLRP3 and DC_LAMP, along with a lower proportion of immunosuppressive subsets Macro_CCL13 and Macro_CST3, in CD47-S dominant samples **(Fig. 2K)**.

**Fig. 2.**
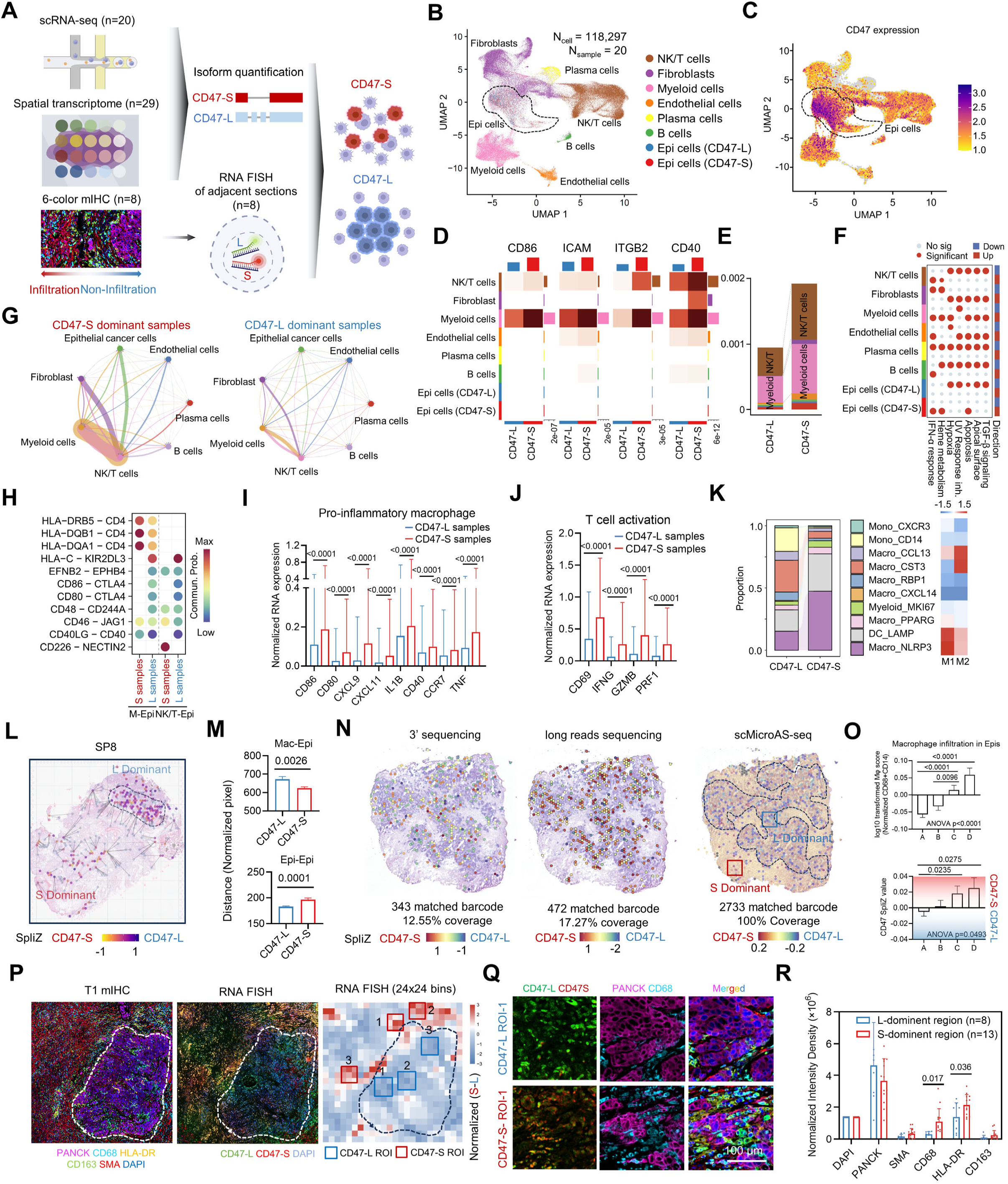
CD47-L, but not CD47-S is preferentially expressed in ovarian cancer cells associated with immunosuppressive tumor microenvironment. **(A)** Flowchart illustrating the CD47 isoform characteristics at the single-cell and spatial levels in relation to cell-cell communication. Spatial isoform characteristics were validated using multicolor fluorescence staining and RNA FISH. **(B)** Uniform manifold approximation and projection (UMAP) embedding of cells from 20 ovarian cancer samples, colored by major cell types. Epithelial cell group was subclustered into CD47-L and CD47-S cells based on the SpliZ value. **(C)** UMAP projection of CD47 expression level. Dashed circles indicate epithelial cell subgroup. **(D)** Heatmap of interaction strength between CD47-L/S epithelial subgroups and other cells via CD86, ICAM, ITGB2, and CD40 signaling. **(E)** Stacked bar plot indicated the global interaction strength of CD47-L and CD47-S subgroups with other cell types. **(F)** Bubble diagram indicating the enriched pathway of CD47-L/S epithelial cell subgroups and other cell types. **(G)** Cell-cell communication network illustrating interactions between myeloid cells, fibroblasts, endothelial cells, NKT cells, B cells, pre-B cells, and epithelial cells in CD47-L/S dominant samples. The thickness of the connections represents the strength of the interactions. **(H)** Dot plot showing the communication probability (Commun. Prob.) of ligand-receptor interactions between M-Epi and NKT-Epi in CD47-S/L dominant samples. The color gradient represents communication strength, ranging from low (blue) to high (red), and dot size indicates interaction significance. **(I, J)** Normalized RNA expression of T cell activation markers **(I)** and pro-inflammatory markers of macrophage **(J)**. Blue, CD47-L epithelial cell subgroup; red, CD47-S epithelial cell subgroup. **(K)** Myeloid subgroups proportions in CD47-L/S dominant samples. Pro-inflammatory M1 score and immunosuppressive M2 score of each myeloid subgroups were annotated on the right. **(L)** Cell communication network of epithelial cells and macrophages cross the SP8 tissue section. Lines mark the nearest tumor cells adjacent to each macrophage. Colors represent SpliZ values in epithelial cells. Dashed lines outline L-dominant epithelial cell region. **(M)** Distance between macrophage and nearest epithelial CD47-L/S spots (up). Distance of the epithelial spot from its nearest neighbor in the CD47-L/S group (down). **(N)** Spatial distribution map of CD47 SpliZ value calculated based on 3’ sequencing, long reads sequencing, and scMicroAS-seq data from same spatial transcriptome cDNA library of OVCST tissue. Each hexagon represents a spatial spot, with color intensity reflecting markers levels. Dashed lines outline epithelial cell region. **(O)** Macrophage infiltration score (Up) and CD47 SpliZ value (Down) in Epithelial cells A, B, C, D subgroups. **(P)** Multiple IHC and RNA FISH image of T1 tissue. Epithelial marker (PANCK, magenta), macrophage markers (CD68, cyan; HLA-DR, yellow; CD163, green), smooth muscle marker (SMA, red), and nuclear stain (DAPI, blue). Heatmap of normalized S-L value determined by florescence intensity of RNA FISH in T1 tissue (24×24 bins). Red square, CD47-S ROI; blue, square, CD47-L ROI (right). Dashed lines outline epithelial cell region. **(Q)** RNA FISH (first column) and immunofluorescence (second and third column) images of corresponding CD47-L ROI-1 and CD47-S ROI-1. **(R)** Normalized intensity density of DAPI, PANCK, SMA, CD68, HLA, and CD163 in CD47-L (blue, n=8) and CD47-S (red, n=13) dominant regions. Intensity density was normalized to DAPI density.

To investigate spatial distribution of CD47-L/S in TME of ovarian cancer, we integrated spatial transcriptome data from 29 cancer samples and identified 7 clusters including 4 epithelial clusters. Strikingly, CD47-L dominant spots located at regions where macrophage-epithelial interaction is lacking, whereas CD47-S dominant spots located at macrophage-infiltrated regions **(Fig. 2L).** Consistently, CD47-L-dominant spots exhibited significantly greater distances from macrophages compared to CD47-S-dominant spots **(Fig. 2M)**. To gain further evidence, we employed spatial transcriptomics by scMicroAS-seq, as well as 3’ sequencing and long-read sequencing on a fresh SOC sample. scMicroAS-seq demonstrated higher spot coverage in detecting CD47 alternative splicing events compared to long-read sequencing (100% vs 17.27%) **(Fig. 2N)**. Consistent with public spatial transcriptome data, SpiZ value (calculated from scMicroAS-seq) was increased along with macrophage infiltration **(Fig. 2O)**, indicating that CD47-S was positively associated with increased infiltration of macrophages in the tumor microenvironment.

We further performed RNA-FISH and multiplexed immunofluorescence staining on adjacent tissue sections from eight tissue samples (T1-T8). The whole tissue was divided in to 24×24 bins and calculated the florescence intensity of CD47-L/S, SMA, PANCK, CD68, HLA-DR and CD163. We found that CD47-L dominant bins located at the boundary of PANCK^+^ regions and normalized CD47-L - CD47-S value was positively correlated with PANCK signal **(Fig. 2P)**. Through analyzing 21 ROIs (Region of Interest), we found CD47-S dominant regions were enriched HLA-DR^+^ (pro-inflammatory) macrophages **(Fig. 2Q-R)**, indicating CD47-S correlates with pro-inflammatory macrophage infiltration within the tumor microenvironment. Together, these results suggest that CD47-L and CD47-S exhibited different distributions in tumor microenvironment. CD47-L but not CD47-S is preferentially expressed in ovarian cancer cells associated with immunosuppressive tumor microenvironment.

### CD47-L and CD47-S display distinct subcellular localization and function

In an attempt to functionalize of CD47-L and CD47-S, we compared the sequences of two isoforms and found CD47-L and CD47-S differ at the intracellular domain, CD47-S is predicted to produce a truncated cytoplasmic tail due to lack of exon 9 and exon 10 (**Fig. 3A)**. To examine the subcellular localization of CD47-L and CD47-S, we constructed GFP-fused CD47-L and mCherry-fused CD47-S and introduced them into OVCAR3, HEY, and HEK293 cells. Confocal 3D images illustrated that mCherry-CD47-S failed to localize to the plasma membrane as GFP-CD47-L did, and instead was retained in Golgi apparatus **(Fig. 3B-C)**. We subsequently quantified mean florescence intensity (MFI) of cell surface and intracellular area by flowmetry in OVCAR3, HEY, and HEK293 cells transfected with GFP-fused CD47-L/S. As a result, CD47-L was higher enriched in cell surface than CD47-S in three cell lines **(Fig. 3D-E)**. Consistently, we observed strong plasma membrane staining in OVCAR3, OVCAR8 cells (high endogenous CD47L/S ratio) and strong intracellular staining in HEY, CAOV3 cells (low endogenous CD47L/S ratio) measured by immunofluorescence staining using anti-CD47 antibody (**Fig. 3F)**. Overexpressing CD47-L, but not CD47-S exhibited strong plasma membrane staining in HEY cells **(Fig. 3G-H)**. We used mass spectrometry to screen for CD47-L isoform-specific interacting proteins and found that RAB32-mediated vesicular transport promoted its distribution to plasma membrane. In addition, the mRNA of the CD47-L isoform was more stable.

**Figure 3.**
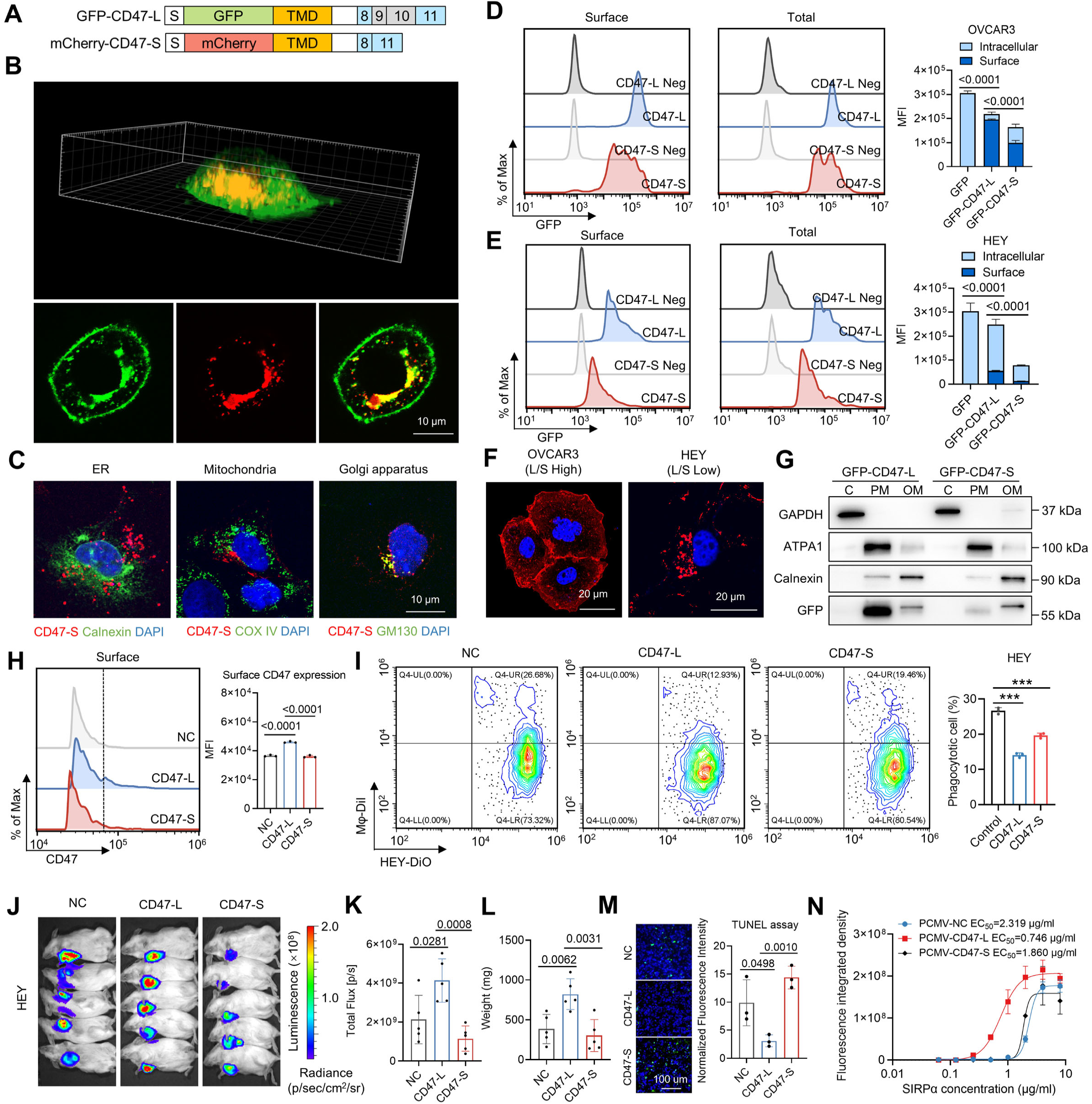
CD47-L and CD47-S display distinct subcellular localization and function. **(A)** Schematic representation indicated the structure of fusion proteins incorporating GFP or mCherry, CD47 transmembrane domain (TMD) and C-terminal variable regions. **(B)** Three-dimensional reconstruction confocal image of OVCAR3 cells expressing GFP fused with CD47-L fragment and mCherry fused with CD47-S. **(C)** Immunofluorescence assay indicated the subcellular location of CD47-S, endoplasmic reticulum (Calnexin), mitochondria (COX IV), and Golgi apparatus (GM130) in OVCAR3. **(D-E)** Mean florescence intensity (MFI) of surface and intracellular CD47 in OVCAR3 **(D)** and HEY **(E)** cells detected by cell flowmetry. Unstained cells are shown in grey. **(F)** Immunofluorescence assay detecting surface and cytoplasmic CD47 expression in OVCAR3 and HEY cells. OVCAR3 was representative CD47-L dominant (L/S high) cell line while HEY was CD47-S dominant (L/S low). **(G)** Plasma and organelle membrane protein isolation assay detecting GFP-CD47-L and GFP-CD47-S subcellular location. C, cytosol; PM, plasma membrane; OM, organelle membrane. **(H)** MFI of membrane CD47 of HEY cells overexpressing CD47-L and CD47-S detected by cell flowmetry. **(I)** Phagocytotic cell percentage of HEY cells overexpressing CD47-L and CD47-S. Dil and Dio double positive cell percentage of HEY overexpressing CD47-L and CD47-S co-cultured with macrophages. **(J)** Luciferase signals of subcutaneous injected immunodeficient mouse and photon flux quantification. Immunodeficient mouse were injected with luciferase-expressing HEY cells with CD47-L or CD47-S overexpression (n = 5). THP1-derived macrophages were replenished through the tail vein every three days from the onset of stable tumor formation. **(K-L)** Total flux **(K)** and tumor weight **(L)** of the xerograph mouse model after CD47-L/S overexpression. **(M)** TUNEL assay detecting dead cells in CD47-L and CD47-S overexpressing xerograph tumor. **(N)** Affinity assay of SIRPα protein with OVCAR3-GFP cells overexpressing CD47-L and CD47-S. The recombined SIRPα protein was immobilized at the bottom of the container and the fluorescence intensity was detected after gentle washing.

Moreover, CD47 protects cancer cells from phagocytosis. Coculture phagocytosis experiments revealed that the CD47-L overexpressed HEY and OVCAR8 cells were less efficiently phagocytosed by macrophages than that of CD47-S overexpressed ovarian cancer cells **(Fig. 3I)**. In mouse xenograft models, CD47-L/S overexpressing HEY cells were subcutaneously injected into immunodeficient mice followed by macrophage supplementation to evaluate their role in ovarian cancer progression. CD47-L overexpression increased the tumor burden and prohibited cell death and macrophage infiltration **(Fig. 3J-M)**. CD47 act by binding and activating ins receptor SIPRα on macrophages^14^. We further performed in vitro binding assays and found that CD47-L had a significantly higher binding affinity for the SIPRα compared to CD47-S **(Fig. 3N)**. Together, these results suggest that two isoforms of CD47 have different subcellular localization and distinct cellular functions. In contrast with CD47-S, CD47L was enriched in cell surface of cancer cells and hence prevent macrophage-mediated phagocytosis by interacting with SIPRα.

### HNRNPA1 regulates isoform switch of CD47-L/S in ovarian cancer cells

To identify splicing factors that regulate the splicing switch between CD47-L and CD47-S, we analyzed the data of a large-scale pooled CRISPRi screen with single-cell RNA sequencing (Perturb-seq, GSE264667, Replogle JM et al.). The library targeting 2,285 essential genes including 780 RNA-binding proteins. Totally 262,956 cells were included and 129,020 cells were found with CD47 SpliZ value. We identified 13 splicing factors that potentially involved in regulating isoform switch of CD47-L/S **(Fig. 4A-C)**. In parallel, an RNA pull-down assay followed by mass spectrometry in HEY cells were performed using an in vitro-transcribed biotin-labeled RNA probe. We identified 38 splicing factors, including multiple hnRNP family members, could potentially bind to CD47 pre-mRNA (**Fig. 4D**). We further obtained 22 splicing factors correlated with CD47 micro-exons skipping level (PSI) and 16 splicing factors correlated with CD47-S percentage based on the TCGA-OV RNA-seq data. Overlap analysis of these three datasets identified HNRNPA1, which was down-regulated in ovarian cancers, as a strong candidate that regulates splicing switch between CD47-L and CD47-S **(Fig. 4E)**. HNRNPA1 was negatively correlated with CD47 exon 9 and 10 inclusion levels in RNA-seq data of TCGA-OV and CCLE ovarian cancer cell lines **(Fig. 4F)**. We validated the splicing switch of CD47 using quantitative RT-PCR and microAS-seq and found that downregulation of HNRNPA1 significantly increased CD47L/S ratio to generate more CD47-L **(Fig. 4G-H)**, whereas overexpression of HNRNPA1 had the opposite effect. We further designed a CD47 dual-fluorescence splicing reporter system that expresses either ZSgreen when and exon 9 and 10 is inclusion or mCherry when exon 9 and 10 in skipped. We observed a large population of cells expressed ZSgreen in cells upon HNRNPA1 knockdown. In contrast, mCherry-positive cell increased in HNRNPA1 overexpressed cells **(Fig. 4I)**. To verify the direct regulation of HNRNPA1 on splicing switch of CD47, we performed an optimized LACE-seq (a method that can identify RNA binding protein targets in low-abundance cells)^15^, called LACE-seq2, in HEY cells. Importantly, HNRNPA1 was shown to bind on CD47 pre-mRNA identified by LACE-seq2, as well as eCLIP-seq and CLIP-seq data **(Fig. 4J-L)**. Consistently, RNA pull-down assay showed that HNRNPA1 was successfully pulled down by biotin-labeled two fragments of CD47 pre-mRNA **(Fig. 4M)**, which was further verified by dot blot hybridization assay **(Fig. 4N).** Moreover, an RNA electrophoretic mobility shift assay (EMSA) confirmed the interaction between HNRNPA1 and CD47 pre-mRNA **(Fig. 4O, P)**. These results suggest that the splicing switch between CD47-L and CD47-S in ovarian cancer cells is regulated by HNRNPA1.

**Figure 4.**
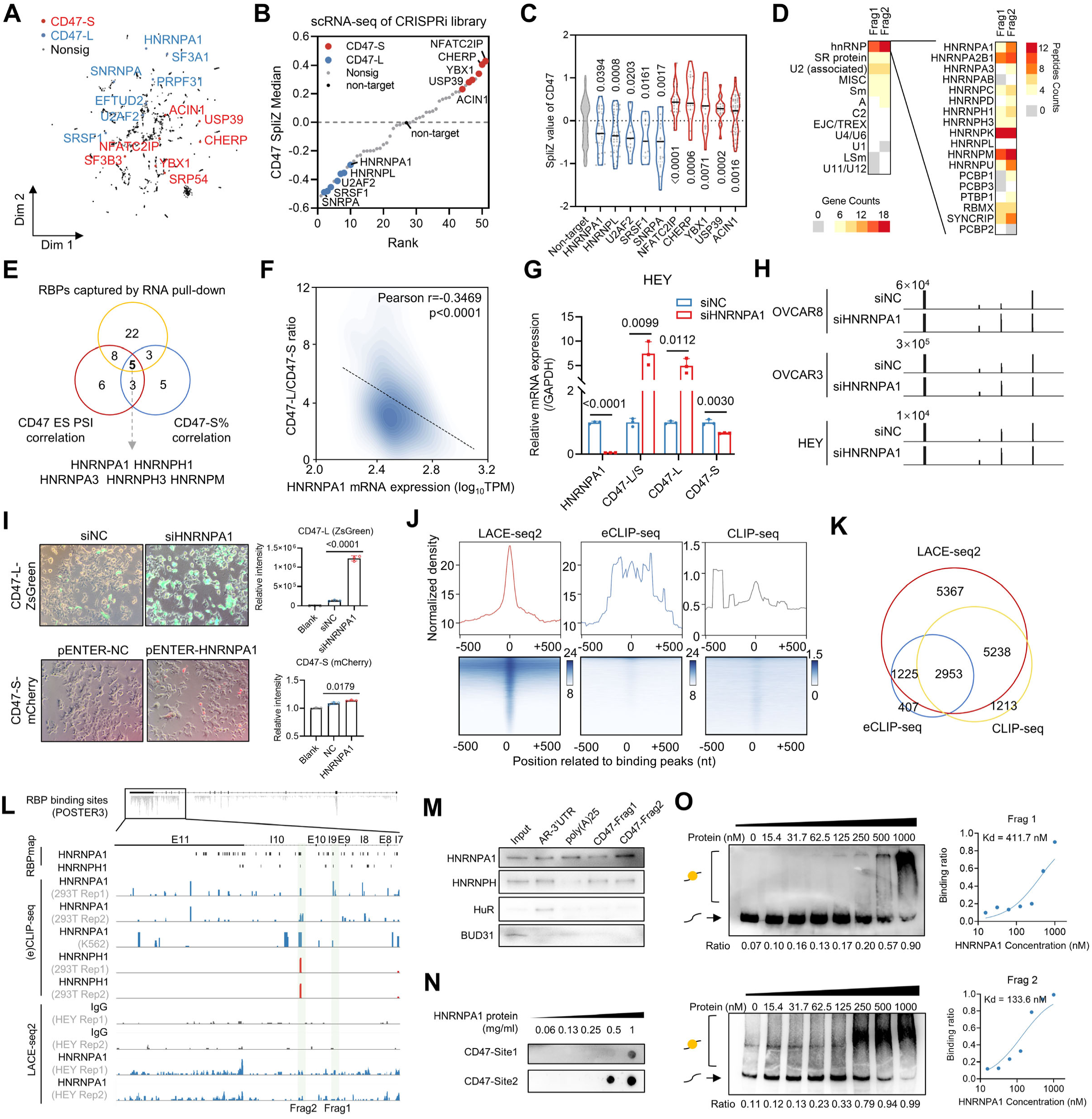
HNRNPA1 regulates isoform switch of CD47-L/S in ovarian cancer cells. **(A)** Minimum distortion embedding of single cell CRISPR data where each dot represents a genetic perturbation. Core splicing factor which induced CD47-L (Blue) and CD47-S (Red) switch tendency are annotated. **(B)** Distribution of medium of CD47 SpliZ value. SpliZ values of cells with core splicing factors perturbation were compared with non-targeted cells. Perturbation induced CD47-L and CD47-S switch were colored in blue and red. **(C)** Violin plot of the cells belonging to each core splicing factor perturbation, comparing to the non-targeted cells. **(D)** RNA pull-down coupled with mass spectrometry screening for CD47 intron interacting RNA binding proteins (RBPs) and their belonging spliceosome complex. **(E)** Venn diagram of the 5 common intron interacting RBPs whose expression was both correlated with CD47 multi-exon skipping PSI and isoform percentage. **(F)** Correlation analysis between CD47 micro-exons skipping PSI and HNRNPA1 mRNA expression. **(G)** mRNA expression levels of HNRNPA1, CD47 and its isoforms after knockdown HNRNPA1 in HEY cells. **(H)** Genomic visualization around CD47 exon 9 and 10 of MicroAS-seq data after HNRNPA1 knockdown in OVCAR8, OVCAR3, and HEY cells. **(I)** Representative images and relative fluorescence intensity quantification of CD47 multi-exon skipping reporter system after HNRNPA1 overexpression and knockdown in HEY cells. **(J)** HNRNPA1 crosslink signal density detected by LACE-seq2, eCLIP-seq, and CLIP-seq related to the binding peaks identified by LACE-seq2. **(K)** Venn diagram of the binding genes identified by LACE-seq2, eCLIP-seq, and CLIP-seq. **(L)** Genomic visualization of CLIP-seq and LACE-seq binding signals of HNRNPA1, HNRNPH1, HNRNPA2B1, HNRNPF, HNRNPM. HNRNPA1 and HNRNPH1 binding sites predicted by RBPmap based on known motifs. Y-axis: 0-10. **(M)** RNA pull-down assay indicated HNRNPA1 and HNRNPH1 interacts with CD47 intron fragment 1 and 2. HuR, which was detected to interact with AR-3’UTR, was the positive control while splicing factor was the negative control. **(N)** Modified dot blot assays were performed with RNA fragments 1 and 2 immobilized onto nylon membranes and their interaction with gradient concentrations of HNRNPA1 was determined. **(O)** RNA EMSA assay detecting the interaction of HNRNPA1 with CD47 intron fragment 1 and 2.

### Isoform switch of CD47 by ASOs provokes macrophage-mediated pyroptosis in ovarian cancer

We next sought to develop antisense oligonucleotides (ASOs) that functions to switch CD47 splicing from CD47-L to CD47-S and suppress tumor growth. We designed 10 ASOs covering HNRNPA1 binding sites adjacent to exon 9 and 10 of CD47 pre-mRNA and transfected into OVCAR3 cells **(Fig. 5A)**. We first examined isoform switch between CD47-L and CD47-S using microAS-seq and observed ASO-I10 and ASO-I9b were two ASOs that significantly induced exon 9 and 10 skipping simultaneously **(Fig. 5B)**. Further qPCR and immunoblotting experiments showed that ASO-I10 was the most potent ASO that decreased CD47-L/CD47-S ratio and CD47-L protein level **(Fig. 5C-D)**. The half-maximal effective concentration (EC50) value for ASO-I10 was 28.97 nM in OVCAR3 cells **(Fig. 5F)**. Dual-fluorescence splicing reporter system indicated the CD47-L/S ratio significantly decreased in 48 h after ASO-I10 transfection **(Fig. 5G)**. Measuring the fluorescence intensity by flow cytometry showed that CD47 levels on the cell surface was significantly decreased after treatment with ASO-I10 **(Fig. 5H)**. Meanwhile, ASO treatment significantly promoted phagocytosis of cancer cells by macrophages illustrated by coculture of macrophages and ovarian cancer cells **(Fig. 5I-J)**, which was further confirmed by 3D spheroid model generated from OVCAR3 and HOC7 cells **(Fig. 5K)**.

**Figure 5.**
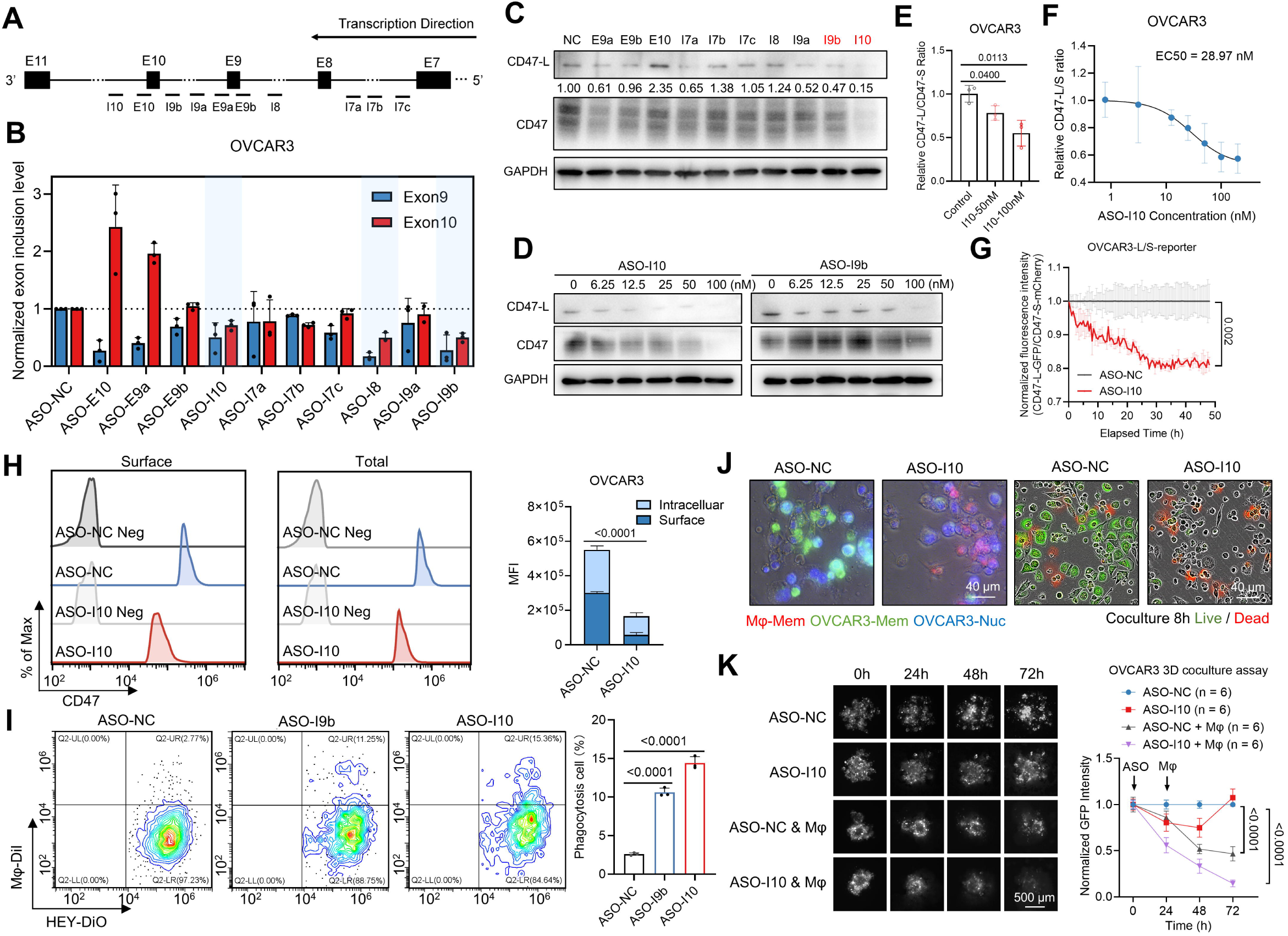
Isoform switch of CD47 by ASOs provokes macrophage-mediated phagocytosis in ovarian cancer. **(A)** Schematic map of the location on the genome of ASOs targeting CD47 multiple exons skipping. **(B)** Normalized exon inclusion level detected by MicroAS-seq after ASO screening in OVCAR3 cells (n=3 biological replicates). **(C)** CD47-L and total CD47 protein expression after ASOs transfection. The bands were quantified and normalized to NC group. **(D)** CD47-L and total CD47 protein expression after transfection with gradient concentrations of ASO-I10. **(E)** Relative CD47-L/CD47-S ratio detected by isoform specific qPCR after 50 nM and 100 nM ASO-I10 transfection. **(F)** Half maximum effective ASO-I10 concentration for inducing CD47-L to CD47-S conversion. **(G)** Normalized fluorescence intensity of OVCAR3-L/S-reporter cell line after ASO-NC, ASO-I10 transfection, detected by long-term imaging of living cells. **(H)** Mean florescence intensity of surface and intracellular CD47 in OVCAR3 cells after ASO-I10 transfection. **(I)** Phagocytosis ratio after ASO-NC, ASO-I9b, and ASO-I10 transfection in OVCAR3 and HEY cells. **(J)** Representative fluorescence images of macrophages (Red) co-cultured with OVCAR3 (Green) superimposed on light microscopy images (Up). The live and dead cells of the co-culture system were also stained with PI (Red) and Calcein-AM (Green) (Down). **(K)** OVCAR3 cells 3D spheroids (n = 6 per group) were co-cultured with macrophages and treated using ASO-I10. Normalized GFP fluorescence intensities were quantified.

Surprisingly, when we monitored time-lapse images of cocultured macrophages and OVCAR3 ovarian cancer cells after treatment with ASO-I10, or HEY cells, we observed macrophages aggregated around tumor cells and thereafter the tumor cells undergo pyroptosis **(Fig. 6A-B)**. The morphological features of ovarian cancer cells undergoing pyroptosis was characterized by light microscope **(Fig. 6C)**, transmission **(Fig. 6D)** and scanning electron microscopy **(Fig. 6E).** Further immunoblotting analysis revealed that the expression levels of pyroptosis-related proteins cleaved-caspase-1 and GSDMD-NT were significantly increased after treatment with ASOs **(Fig. 7G)**. Increased secretion of IL-1β and IL-18 was observed in medium of cocultured macrophages and OVCAR3 or HOC7 cells after treatment with ASOs measured by Enzyme-linked immunosorbent assay (ELISA) **(Fig. 6H-I)**. Immunofluorescence co-staining further showed the colocalization of GSDMD in ovarian cancer cells, but not in macrophages **(Fig. 6F)**. We next performed single-cell RNA sequencing on cocultured macrophages and OVCAR3 cells after treatment with I10 or negative control. As expected, the spliZ value was significantly increased after treatment with ASO-I10, indicating ASOs effectively induces isoform switch from CD47-L to CD47-S **(Fig. 6J)**. Importantly, we observed a less proportion of epithelial-cancer cells in cocultured macrophages and OVCAR3 cells treated with ASO-I10 compared to negative control **(Fig. 6K-L)**. Pseudotime analysis of macrophages subclusters revealed that ASOs treatment promoted macrophages transform into C1QC^+^ macrophages with enhanced phagocytotic activity and adhesion (**Fig. 6M-N)**. Besides, ASO-I10 treated epithelial-cancer cells showed higher expression of pyroptosis related response genes. These results indicate that ASOs targeting isoform switch from CD47-L to CD47-S stimulate macrophage-mediated phagocytosis of ovarian cancer cells.

**Figure 6.**
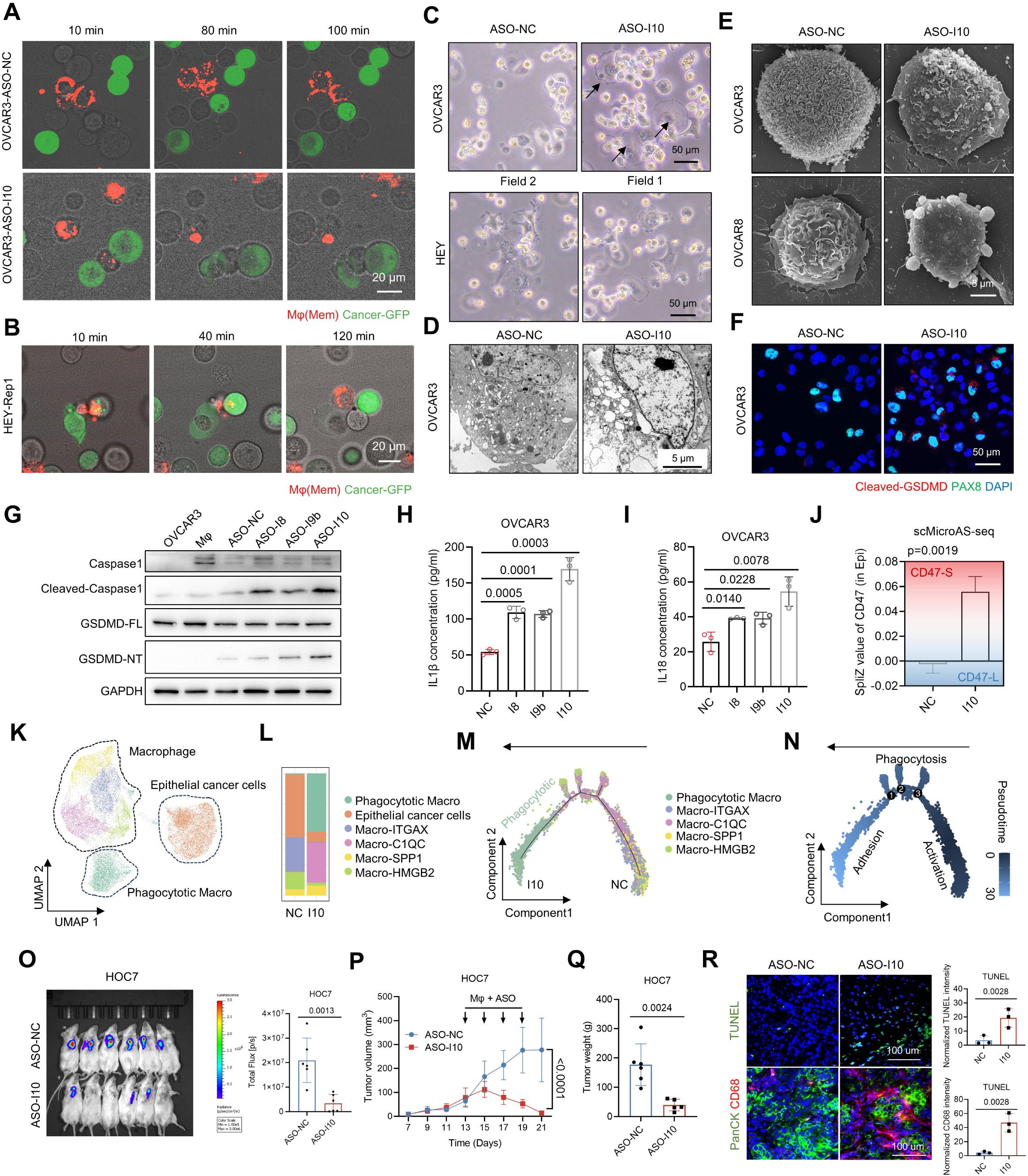
ASO promotes macrophage induced cancer cell proptosis and inhibits tumor progression. **(A)** Time-lapse photographs of ASO-NC and ASO-I10-transfected OVCAR3-GFP in co-culture with Dil-stained macrophages within 100 mins. **(B)** Time-lapse photographs of HEY-GFP (CD47-L/S low) in co-culture with Dil-stained macrophages within 120 mins. **(C-D)** Representative photographs of HEY or ASO-transfected OVCAR3 cancer cells co-cultured with macrophages captured by light microscopy **(C)** and transmission electron microscopy **(D)**. **(E)** Representative images of scanning electron microscopy of ASO-I10 transfected OVCAR3 and OVCAR8 cells after co-culture with macrophages. **(F)** Distribution of PAX8 (Green) and Cleaved-GSDMD (Red) in ASO-transfected OVCAR3 cells co-cultured with macrophages was determined using immunofluorescence. **(G)** Protein expression of pyroptosis markers after ASO transfected OVCAR3 cells co-cultured with macrophages. **(H, I)** IL18 and IL1beta concentration in supernatants of ASO-transfected OVCAR3 cells co-cultured with macrophages. **(J)** CD47 SpliZ index in scRNA-seq data from a macrophage and cancer cell co-culture system following ASO-I10 treatment, as detected by targeted amplification using scMicroAS-seq. **(K)** UMAP projection of cell clusters in scRNA-seq of macrophage co-cultured with ASO-I10 transfected OVCAR3 cells. **(L)** Proportions of epithelial cancer cells and macrophages clusters with ASO-NC and ASO-I10 transfection. **(M)** Pseudotime trajectory of the macrophage cluster co-cultured with ASO-NC and ASO-I10 transfected OVCAR3 cells, colored by macrophage clusters. **(N)** Pseudotime trajectory of the macrophage cluster within the co-culture system. The enriched pathways of marker genes of each stage were annotated. **(O)** Luciferase signals of subcutaneous injected immunodeficient mouse and photon flux quantification. Immunodeficient mouse were injected with luciferase-expressing HOC7 cells mixed with primary human fibroblast (n = 6). THP1-derived macrophages were supplemented through the tail vein after stable tumor formation. LNP packaged ASO was delivery through intratumor injection from Day13. **(P-Q)** Tumor volume and weight of the xerograph mouse model after ASO treatment and macrophage supplement starting from Day 13. **(R)** Immunofluorescence and TUNEL images of xenograft tumor model of ASO-NC and ASO-I10 groups. PANCK (green) represents epithelial tumor cells and CD68 (red) represents macrophages.

Next, we investigated the anti-tumor effect of ASOs in vivo. A cell-derived xenograft model was established using luciferase-expressing HOC7 cells and primary cancer associated fibroblast, exhibiting histological features similar to high-grade serous ovarian carcinoma (HGSOC). The anti-tumor effect was assessed in M-NSG mice replenishing with THP1-derived macrophages. ASOs treatment significantly promoted pro-inflammatory infiltration and resulted in a low tumor burden (**Fig. 6O-R)**. Consistently, in BALB/c nude mice of xenografted OVCAR8 ovarian cancer cells, ASOs treatment also showed significant anti-tumor activity compared to control. The anti-tumor activity of ASOs was further confirmed in a patient-derived xenograft models of ovarian cancer. We had also provided evidences that ASOs had no detectable immunogenicity and cytotoxicity on macrophages, less hemolytic effects in contrast to CD47 antibody. Collectively, these data illustrate that ASOs that switch CD47 splicing from CD47-L to CD47-S exhibit significant antitumor effects.

## Discussion

This study provides novel insights into the role of CD47 isoform switching in the immune tumor microenvironment and its implications for immunotherapy of human malignancies including ovarian cancer. Our analysis on long read RNA sequencing data revealed that two micro-exons (exon 9 and exon 10) of CD47 were alternative spliced to produced two isoforms with distinct subcellular localization and function. Normal tissues preferentially express CD47-S (lacking exon 9 and exon 10), that was unable to locate at the cell surface to bind with SIRPα as CD47-L did. In contrast, ovarian cancer tissues exclusively express CD47-L that incudes exon 9 and exon 10, that localize to the plasma membrane and bind with SIRPα to trigger “don’t-eat-me” signal. Importantly, the high level of CD47-L is associated with worse prognosis in ovarian cancer patients. Human CD47 gene has 13 exons and four alternative spliced isoforms of CD47 mRNA have been discovered^16^. In line with our findings, CD47-S (isoform 2) is the most abundant isoform in normal tissues such as endothelial and epithelial cells^17^. The functional significance of spliced CD47 is largely unknown. Our results functionally characterized the two isoforms of CD47, which was regulated by splicing factor HNRNPA1. Isoform switch from CD47-L to CD47-S implying a promising approach to improve immune responses against cancers.

Our work further characterized the expression pattern of CD47-L and CD47-S in TME of ovarian cancer by analyzing scRNA-seq and spatial transcriptome data. Cell-cell communication analysis on 20 ovarian cancer samples of scRNA-seq data revealed that CD47-S dominant samples had stronger communication strength with immune-active tumor microenvironment compared to CD47-L dominant samples. Importantly, spatial transcriptome data showed that CD47-L dominant spots located at regions where macrophage-epithelial interaction is lacking, whereas CD47-S dominant spots located at regions where macrophage-epithelial interaction is active. These data suggest that CD47-L, but not CD47-S contributes immunosuppressive tumor microenvironment.

Human CD47-blocking antibodies have demonstrated promising results in preclinical studies and phase 1/2 clinical trials in solid tumors including lung^18^, breast^19^, pancreatic^20^, colorectal^14^, and ovarian cancers^21^. CD47-blocking therapies could enhance the efficacy of other immune checkpoint inhibitors^22^. However, CD47 blockade can cause anemia and other hematological toxicities because CD47 is highly expressed on red blood cells and platelets. Splice-switching ASOs have been approved by FDA for the treatment of spinal muscular atrophy (SMA)^23^. Here, we developed ASOs that successfully switch CD47 splicing from CD47-L to CD47-S, hence decreased CD47 protein level. Our data further demonstrated that ASOs targeting isoform switch of CD47-L to CD47-S exhibited effective anti-tumor activity in both in vitro and in vivo. In contrast to CD47 blocking antibodies, ASOs showed less toxicity on red blood cells since ASOs target CD47 at the pre-mRNA level in the nucleus of the cell. Therefore, we provide a novel therapeutic strategy targeting CD47-SIRPα immune checkpoint.

Of note, our data further suggest a mechanism that ASOs treatment stimulated macrophage phagocytic activity and further triggers pyroptosis in ovarian cancer cells. In addition to phagocytosis, CD47 blockade exert their anti-tumor effect through various mechanisms^24^. Anti-CD47 antibody treatments have been shown to activate CD8^+^ T-cells and dendritic cells^25^, enhance NK-cell mediated cytotoxicity^26^. CD47 inhibition can also induce tumor cell apoptosis through reduce the expression of anti-apoptotic genes^27^. Triggering pyroptosis in cancer cells is emerging as a promising strategy for cancer therapy^28^. Our data provide strong evidence that Splice-switching ASOs targeting CD47 trigger macrophage-mediated pyroptosis dependent on caspase-1 and GSDMD, suggesting macrophage-mediated tumor cell pyroptosis as a key anti-tumor mechanism targeting on CD47.

In summary, our work screened immune checkpoints that could be regulated by alternative splicing and identified CD47 can generate two major isoforms that differentially expressed in normal and ovarian cancer tissues. CD47-L, but not CD47-S is preferentially expressed in ovarian cancer cells associated with immunosuppressive tumor microenvironment. Notably, splice-switching ASOs targeting CD47L/S evoked macrophage-mediated antitumor immune response and inhibited tumor progression. Overall, our findings highlight the potential of splicing modulators in immune checkpoint blockade therapy.

### Limitations of the study

There are several limitations remain to be resolved. Firstly, subcutaneous CDX and PDX tumor models were used to test anti-tumor effect of ASOs targeting isoform switch of CD47. Humanized mouse models that mimic the human tumor microenvironment and immune response, would further support our findings. Secondly, cell-cell communication analysis on scRNA-seq data revealed that CD47-S^high^ epithelial cancer cells interacted more closely with NK/T cells and myeloid cells compared with CD47-L^high^ epithelial cancer cells, supporting the notion that CD47 inhibition can enhance NK/T cells response^10^. Further investigation would be required to understand the effect of CD47 on NK/T cells as well as macrophage. Additionally, we found that splice-switching ASOs targeting CD47 trigger macrophage-mediated pyroptosis, however, the exact mechanism underlying this process remain unclear further investigations are needed.

## Methods

### Human tissue samples

SOC specimens and FTs were collected from April 2005 to January 2023 in Qilu Hospital. The SOC samples were obtained from patients with primary ovarian cancer who had not undergone any previous surgery or chemotherapy. In addition, FTs were obtained from patients who underwent total hysterectomy and bilateral salpingo-oophorectomy for uterine diseases or benign neoplastic adnexal pathological changes. Fresh tissue samples were collected within 2 h of surgery and were sliced to 5 mm^3^ and immersed in 10 vol of RNALater (Ambion, Austin,TX). The tissue samples were stored at –80 °C. All patients provided informed consent, and ethical approval was granted by the Ethics Committee of Qilu Hospital, Shandong University (KYLL-202311-021).

### Cell line and cell culture

HEK293, HEK293T, NIH-OVCAR3, OV90, THP-1 and CAOV3 were purchased from ATCC (American Type Culture Collection). FTE187, HEY, A2780 were obtained from the Jian-Jun Wei lab, Northwestern University. OVCAR8, HOC7 were gifts from the Ding Ma’s lab, Huazhong University of Science and Technology. A2780, HEY, OV90, CAOV3, HEK293 and HEK293T were cultured in high glucose Dulbecco’s modified Eagle’s medium (DMEM) (Macgene). FTE187 cells were cultured in medium consisting of 1:1 Medium199 and MCDB105 medium (Sigma-Aldrich). OVCAR8 and HOC7 and were cultured in RPMI 1640 (Macgene). The medium was supplemented with 10% fetal bovine serum (Gibco) and 1% penicillin/streptomycin (Macgene). THP-1 was cultured in RPMI 1640 medium supplemented with 0.05mM β-mercaptoethanol. All cell lines were validated by STR profiling and tested for mycoplasma contamination. The cells were cultured at 37 °C in a humidified incubator containing 5% CO_2_.

### Cell-derived and patient-derived xenograft model

In the subcutaneous xenograft model detecting CD47 isoforms functions, female M-NSG immunodeficiency mice (NOD-*Prkdc*^scid^*Il2rg*^em1^/Smoc) were obtained from Shanghai Model Organisms Center. Mice were randomly divided into 3 groups (5 mice per group). CD47-L/S overexpressing luciferase^+^ HEY cells (5 × 10^6^ per mice) were mixed with primary cancer-associate fibroblast at 1:1 ratio. The cell suspension was mixed with Plurigel Matrix (GL101, Vazyme) at a 1:1 ratio on ice (4°C) and gently homogenized by pipetting before subcutaneous injection. Three weeks later, the mice were anesthetized with Avertin (20 μl/g body weight, DW3106, Dowobio) and D-Luciferin potassium salt (15 mg/ml, dissolved in DPBS, 150 μg/g body weight, 50227, Sigma) was injected intraperitoneally. Bioluminescence images were captured 10 min later using IVIS Spectrum in vivo imaging system (PerkinElmer).

In the xenograft model detecting ASOs anti-tumor function, HOC7, OVCAR8 cell line-derived xenograft and patient derived xenograft were used. In the living image HOC7 xenograft model, female M-NSG immunodeficiency mice (Shanghai Model Organisms Center) were subcutaneously injected with 100 μl of a mixture of HOC7 and primary fibroblast cells resuspended in Plurigel Matrix. When the subcutaneous tumors reached a volume of 100 mm^3^ at day 13, mice were randomly divided into two groups (6 mice per group). 12.5 μg ASO per mice were delivered with CALNP RNA in vivo (DN004, D-Nano Therapeutics) lipid nanoparticle system. 12.5 μl ASO (1 mg/ml, dissolved in nuclease-free water) were mixed with 25 μl Reagent A. Add 100 μL of Reagent B, mix thoroughly by pipetting more than 20 times, and incubate at room temperature for 10 minutes, following intratumor injection. 3×10^6^ THP1-derived human macrophages are replenished by tail vein injection. ASO and macrophages were administered alternately every other day. The in vivo imaging procedure was performed using the same method as described above. OVCAR8 cell line-derived xenograft model was established in female BALB/c-nude mice with functionally intact macrophages (6–8 weeks old, CAnN.Cg-*Foxn1^nu^*/Crl, Vital River Laboratory Animal Technology, 10 mice per group). LNP packaged ASO was intratumoral injected at Day 11 and Day 13 with no additional macrophage replenishment.

To establish the PDX model, fresh tumor tissues were obtained from patients with primary ovarian cancer who had not undergone any previous surgery or chemotherapy. The tissues were immediately transported in cold, sterile solution of sodium chloride (0.9% NaCl) supplemented with antibiotics. Under aseptic conditions, the tumor tissues were minced into small fragments (approximately 2–3 mm³) and subcutaneously implanted into the flanks of 6–8-week-old female M-NSG mice (Shanghai Model Organisms Center). Upon reaching a volume of 1,000 – 1,500 mm³, tumors were harvested and passaged into subsequent generations of mice, and cryopreserved for future use. Histopathological and molecular analyses confirmed the preservation of the original tumor characteristics across passages. Mice with PDX were randomly divided into two groups before treatment (7 mice per group). LNP packaged ASO and macrophages were administered every other day from Day 10. Tumors were harvest when maximum volume reaching 1,500 mm³.

Mice were monitored for tumor growth every other day, and tumor volume was calculated using the formula: (length × width²)/2. At the endpoint of the study, the mice were anesthetized and tumor tissues were excised. Tumor weight was measured, also evaluated macrophage infiltration status through immunofluorescence. Dead cells signal were detected through TUNEL assay according to the manufacturer’s protocol (KEYGEN, KGA703). The maximal tumor diameter permitted is 15 mm, and this limit was not exceeded during the experiments. Less than 5 mice were housed per cage under controlled conditions (20–25 °C, 50% humidity, and a 12-hour light/dark cycle). The Ethical Committee of School of Basic Medical Sciences, Shandong University approved the animal experiment procedures (ECSBMSSDU2021-2-91).

### cDNA cloning, plasmid construction, and transfection

GFP-CD47-L, a plasmid coding GFP tagged CD47 with its own short 3’UTR, were purchased from Addgene (GFP_CD47_SU, #65473). GFP-CD47-S was cloned by deleting exon 9 and exon 10 from GFP_CD47_SU using Mut Express II Fast Mutagenesis Kit V2 (Vazyme). mCherry-CD47-S was constructed by replacing GFP with mCherry using KpnI-EcoRI cloning sites (Tsingke). The full length CD47-L and CD47-S ORFs were cloned from human placenta cDNA and constructed into pLenti-C-Myc-DDK-IRES-Puro lentiviral gene expression vector using AsisI-Mlul cloning sites. PCR reaction was performed using Phanta Max Super-Fidelity DNA Polymerase (Vazyme) and cloning reaction using ClonExpress II One Step Cloning Kit (Vazyme).

For the CD47-L/S splicing reporter system, two minigene reporter plasmids were constructed separately and transfected into the same cell after antibiotic screening. Sequence of exon 8, part of intron 8, exon 9, exon 10, part of intron 10, exon 11 and ZsGreen were synthesized and constructed in to pLVX-IRES-puro using EcoRI-XbaI (Tsingke). Exon 9 and 10 inclusion and intron remove leads to coding form ZsGreen. Sequence of exon 8, part of intron 8, exon 9, exon 10, part of intron 10, exon 11 and mCherry were synthesized and constructed in to pLVX-IRES-neo using EcoRI-XbaI (Tsingke). The remained mini-intron included ∼100 bp near the 3’ and 5’ splicing site and splice factor binding enriched regions. Exon 9 and 10 skipping and intron remove leads to coding form mCherry based on coding shift strategy. The lentivirus vectors were transfected into HEK293T cells together with psPAX2 and pMD2.G to produce lentivirus particles. Stable cell lines were established by lentivirus infection followed by puromycin (2 μg/ml) or blasticidin (15 μg/ml) selection for 2 weeks.

The HNRNPA1-pENTER overexpression plasmid was achieved from WZ Biosciences (CH877838). The DsRed2-ER-5 was purchased from Addgene (#55836). Small interfering RNA (siRNA) of RACK1, RAB32, TCP1, and RPS2 was synthesized by Tsingke. genOFF st-h-HNRNPA1_003 was purchased from RiboBio (stB0006663C-1-5). ASO was synthesized with phosphorothioate modifications in the 5’ and 3’ nucleobases to enhance the stability and binding affinity. Plasmids, siRNAs and ASOs were transfected into cells using jetPRIME transfection reagent (101000046, PolyPlus) following the manufacturer’s instructions. RNA and protein were extracted 48 h or 72 h after transfection.

### RNA isolation and PCR analysis

Total RNA was extracted from cultured cells or fresh tissues with FastPure Cell/Tissue Total RNA Isolation Kit (RC101-01, Vazyme). Then RNA was reverse transcribed into cDNAs using HiScript II Q RT SuperMix for qPCR (R222-01, Vazyme), and qPCR was performed with ChamQ SYBR Color qPCR Master Mix (Low ROX Premixed) (Q431-03, Vazyme) in the Applied Biosystems QuantStudio 3. GAPDH served as the endogenous control. The 2−ΔΔCT method was used for the relative quantification of the qPCR data.

### Western blot

The samples were lysed on ice in Western and IP Cell Lysis Buffer (Beyotime) mixed with 1× Protease Inhibitor Cocktail (TargetMol), and the protein concentration was determined using the bicinchoninic acid protein assay (Beyotime). SDS-PAGE was performed to separate protein samples, which were then electro-transferred onto a PVDF membrane at 200 mA for 90 min. The membrane was blocked with 5% skim milk for an hour at room temperature and incubated over-night with primary antibodies at 4°C. Primary anti-bodies were diluted at a ratio of 1:1000 with Primary Antibody Dilution Buffer (Beyotime) except anti-GAPDH Ab which was diluted at 1:5,000. Horseradish peroxidase (HRP)-conjugated secondary antibodies (1:10,000 dilution, Jackson ImmunoResearch) were incubated for 1h at room temperature. The Enhanced Chemiluminescent detection (ECL) mixture (Vazyme, E411) was evenly applied to the membrane. The chemiluminescent signal was captured using a chemiluminescence imaging system (Tanon 5200). Band intensity was quantified with the ImageJ software (version 1.53c).

### BMDM and THP-1 derived macrophage induction

BMDM was flushed from femurs and tibias of wild-type mice with PBS and filtered through a 70-μm filter. Cells were centrifuged at 1000 rpm for 5 minutes and lysis with 1×RBC Lysis Buffer (BioLegend, 420301) for 10 minutes. Primary macrophage differentiation was induced by 20 ng/mL of murine macrophage colony-stimulating factor (Abclonal, RP01216S) for 7 days in complete RPMI 1640 medium. Induction was validated using PE anti-mouse F4/80 Recombinant Antibody (BioLegend, 157303) by flow cytometry. THP-1 was induced in complete 1640 medium with 100 ng/ml Phorbol 12-myristate 13-acetate (PMA, Selleck, S7791) for 24 hours and removed PMA for at least 24 hours. Induction was validated using APC/Cyanine7 anti-human CD14 antibody (BioLegend, 325619) by flow cytometry.

### Phagocytosis Assay

After transfection 48 hours, both cancer cell and macrophage were stained by DiO or Dil separately (Beyotime, Catalog No. C1419M, C1991M) according to the manufacture’s instruction. The dilution ratio of DiO and Dil were 1:1000 and the staining enhancer was diluted at 1:400. The staining working solution was added to the cell culture plate and incubated at 37°C for 15 minutes. After staining, the cells were washed three times and directly co-cultured (macrophage: cancer cell = 2:1) in Nunclon Sphera 96 well plate (ThermoFisher, 174925). Phagocytotic cell ratio was detected by CYTOFLEX S (Beckman), DiO was detected using the FITC channel, and Dil was detected using the PE channel. Double-positive cells were considered phagocytic cells.

### Scanning and transmission electron microscopy

For scanning electron microscopy, 2×10^4^ cancer cells were seeding on 24-well plate cell culture slides. 2×10^4^ macrophages were added to the well and cocultured for 4-6 hours. Pyroptosis of cancer cell was validated under the light microscope. The samples were gently washed with PBS, fixed with 2.5% glutaraldehyde, and post-fixed with 1% osmium tetroxide for 1-2 hours, followed by dehydration. Samples were then critical-point dried using liquid COL and sputter-coated with a 5-nm layer of gold-palladium to enhance conductivity. Image acquisition of the sample was performed using a JSM-IT700HR (JEOL) scanning electron microscope. For transmission electron microscopy, 3.5 × 10^5^ cancer cells and 3.5 × 10^5^ macrophages were cocultured for 4-6 hours and collected in PBS. Samples were fixed and dehydration with the same protocol. The dehydrated samples were sequentially infiltrated with acetone-Epon-812 mixtures (3:1, 1:1, 1:3), embedded in pure Epon-812 resin, and polymerized at 60°C for 48 hours. Ultrathin sections (60–90 nm) were collected on copper grids, and stained with uranyl acetate (10–15 min) and lead citrate (1–2 min) at room temperature for contrast enhancement. Image acquisition of the sample was performed using a JEM-1400FLASH (JEOL) scanning electron microscope.

### IP-MS

OVCAR3 cells overexpressing GFP-fused CD47 isoforms were harvested, and cell pellets were lysed on ice for 30 min with Western and IP Cell Lysis Buffer (Beyotime). Whole-cell extracts were incubated with 5 μg anti-GFP antibody (ABclonal) overnight at 4 °C on a tube rotator to ensure continuous mixing, then the immunocomplex were captured with magnetic Protein A/G beads (MCE) for 4 h at 4 °C. Beads were washed with Western and IP Cell Lysis Buffer for five times, and the immunocomplex was resuspended in 1× SDS-PAGE loading buffer. The eluted proteins were separated by SDS-PAGE following Coomassie brilliant blue staining. After trypsin digestion and desalting, LC-MS/MS was performed using a Q Exactive plus mass spectrometer (Thermo Fisher Scientific) coupled with an Easy nLC by Genechem. The interacting proteins were also detected by western blot using VeriBlot for IP Detection Reagent (HRP) (Abcam, ab131366).

### Immunohistochemistry staining of tissue sections

Formalin-fixed, paraffin-embedded tissues or tissue microarray sections were initially deparaffinized using xylene and subsequently rehydrated through a graded ethanol series. Antigen retrieval was conducted by heating the samples in EDTA buffer within a microwave. Following this, tissue slides were blocked with 1.5% normal goat serum and then incubated overnight at 4 ° C with primary antibodies anti-CD47 (1:200 dilution, Santa Cruz Biotechnology, sc-12730), anti-HNRNPA1 (1:200 dilution, Abways, CY6867) and anti-HNRNPH (1:250 dilution, Abways, AY9553). After incubation, the sections were treated with a secondary antibody and developed using diaminobenzidine (DAB) for visualization. The immunohistochemical (IHC) staining scores for the tissue microarray, which assessed the staining intensity and proportion across the sections, were quantified by two independent pathologists.

### Immunofluorescence and Multiplex Immunofluorescence Staining

For immunofluorescence, cells on glass coverslips were fixed with 4% paraformaldehyde for 15 min at room temperature followed by permeabilization with 0.1% Triton X-100 in PBS for 10 min at room temperature. Tissue slides were processed as described above for immunohistochemistry staining. Samples were then blocked with 5% normal goat serum in PBS for an hour at room temperature and incubated with primary antibodies overnight at 4 °C and then incubated with fluorophore-conjugated secondary antibody (1:500 dilution) for an hour at room temperature in the dark. 4’,6-diamidino-2-phenylindole (DAPI, ab104139, Sigma-Aldrich) was used to stain cell nuclei. The images were captured on a high-speed confocal platform (Andor, Dragonfly 200) or Olympus BX53 microscope system. TUNEL assay detecting DNA damages was performed using the TUNEL Cell Apoptosis Detection Kit (KEYGEN, KGA703) according to the manufacturer’s protocol. Normalized CD68 and TUNEL intensity were calculated by: raw intensity density / DAPI area. Multiplex immunofluorescence staining was performed using the PANO 6-plex IHC kit (cat. 10081100100, Panovue). Primary antibodies were applied and incubated at room temperature for 30 min sequentially, followed by incubation with HRP-conjugated secondary antibodies and tyramide signal amplification (TSA). After each TSA step, the slides were subjected to microwave treatment. Following the labeling of all human antigens, nuclei were counterstained with DAPI (ab104139, Sigma). Whole-slide fluorescent images were acquired using an TissueFaxs Spectra system (TissueGnostics) equipped with a 20× objective lens and analyzed using QuPath (v0.5.1) and ImageJ (v1.53c). To compare the fluorescence intensity in L- and S- dominant regions, the intensity value was normalized to the relative DAPI ratio in each region.

### Affinity assay of SIRP**α** protein with cancer cells

Diluting the Recombinant Human SIRP-alpha/CD172a Protein (RP00171, Abclonal) into gradient concentration (0.0625-8 ug/ml) with Coating Buffer (RM01756, Abclonal). Coat each well of the Uncoated Plate (RM01758, Abclonal) with 100 µL diluted solution and incubate overnight at 4°C. Aspirate the liquid from each well and wash once by adding 200 µL of Wash Buffer (1×) (RM00026, Abclonal) per well. Block each well with 200 µL of Blocking Buffer (RM01757, Abclonal) for 1 hour at room temperature. After removing residual liquid, 2×10^4^ OVCAR3-GFP cells in culture medium were added to each well and incubate for 30 mins at room temperature. The fluorescence intensity was detected with and quantified with ImageJ after washing three times.

### RNA FISH

RNA FISH was performed using Fluorescence in Situ Hybridization Kit for RNA (Beyotime, R0306M). Tissue section (4 μm) and TMA were dewaxed and rehydration as described above for immunohistochemistry staining. Then it was treated with 10 μg/ml Proteinase K in PBS at room temperature for 5 min. After permeabilization with 0.5% Triton X-100 in PBS for 5 min, tissue was re-fixed with 4% paraformaldehyde for 10 min and washed again. Samples were immersed in 0.5 M HCl for 5 minutes, washed twice with PBS, and treated with freshly prepared Acetylation Solution (Acetylation Solution A : Solution B = 1:400) for 10 minutes at room temperature. Afterward, samples were washed twice with PBS for 5 minutes each. Pre-hybridization was performed using Hybridization Solution with 1×Yeast RNA at 55°C for 20 minutes. Probes were prepared at 100 μM in pre-treated 1×Hybridization Solution and hybridized with sections at 55°C for 12 hours. Post-hybridization washes were performed sequentially with preheated Wash Buffers I (3 washes), II (1 wash), and III (1 wash), for 5 minutes each. Samples were stained with 1 μg/ml DAPI for 3 minutes in the dark, washed twice with PBS, and mounted with Mounting Medium, antifading (Solarbio, S2100). Fluorescence whole-slide scanning was performed using MIDI:3Dhistech (Pannoramic) or TissueFaxs Spectra (TissueGnostics) and analyzed with CaseViewer (v2.4) or TissueFAXS Viewer (v7.1.6245.14). The florescence intensity was further analyzed using ImageJ. The spatial intensity was transformed into test image and divided into 24×24 bins (8-bit). Red-green fluorescence intensity differences were Z-score normalized within each section and visualized with pheatmap (v1.0.12).

### RNA Pull-Down Assay, EMSA, and Dot Blot Analysis

CD47 pre-mRNA fragments were PCR-amplified from human placental genomic DNA using primers containing 5’ T7 promoter sequence. Using the T7 High Yield RNA Transcription Kit (TR101, Vazyme), biotinylated RNA probes were synthesized in vitro and subsequently labeled with the Pierce RNA 3’ End Desthiobiotinylation Kit (Thermo Fisher Scientific). RNA pull-down assays were performed using the Magnetic RNA-Protein Pull-Down Kit (Thermo Fisher Scientific), with poly(A)LL RNA and the 3′ untranslated region of androgen receptor RNA serving as negative and positive controls, respectively. Captured proteins were identified by mass spectrometry (Genechem) and validated via Western Blot.

For electrophoretic mobility shift assays (EMSA), 10 ng of biotin-labeled RNA probe (heat-denatured at 85°C for 3 min to eliminate secondary structures) was incubated with gradient concentrations of recombinant His-tagged HNRNPA1 (CSB-EP010600HU, Cusabio) for 20 min at 25 °C. Using Chemiluminescent RNA EMSA Kit (GS606, Beyotime), RNA-protein complexes were resolved on 5% non-denaturing polyacrylamide gels in 0.25× TBE buffer (120 V, 60 min) and transferred to membranes in 0.5× TBE (390 mA, 40 min). After UV crosslinking (600 mJ, CL-1000 linker), membranes were probed with HRP-conjugated streptavidin, and signals were detected using an ECL system (Tanon 5200). Binding affinity was quantified as the ratio of complex band intensity to the total signal (complex + free probe).

Classical dot blot protocol was modified to detect RNA and protein interaction. Briefly, Gradient concentrations of HNRNPA1 protein were spotted onto a nylon membrane and air-dried. The membrane was then incubated at 37°C for 30 min. Blocking was performed with 10 mL blocking buffer (GS606, Beyotime) for 1 h at room temperature with gentle shaking. After removing the blocking solution, the membrane was incubated with biotin-labeled probe (final concentration: 1 nM) for 1 h at room temperature. The membrane was washed three times with 0.1%Tween-20 in 1 ×TBS buffer (5 min per wash) and UV-crosslinked twice at 350 mJ using a CL-1000 Ultraviolet Crosslinker. Subsequently, the membrane was incubated with diluted HRP-conjugated streptavidin for 1 h at room temperature, followed by three additional washes. Finally, signals were detected using ECL substrate and imaged.

### Flow Cytometry for Surface and Intracellular Protein Detection

Surface and intracellular protein was detected according to previous study.^29^ For surface CD47 detection, the cells were harvested and incubated with mouse anti-CD47-PerCy5.5 (BD Biosciences, 561261) for 30 min at 4L in FACS Buffer A (PBS containing 0.5% FBS). For detection of surface GFP, cells were incubated with Rabbit anti GFP-Tag mAb (Abclonal, AE078), then incubated with goat anti-rabbit Alexa Fluor 594 (Invitrogen, A11012) for 30 minutes at 4°C. The cells washed three times with FACS Buffer A before analyzed on Cytoflex S (Beckman). For intracellular protein detection (to detect total protein expression), cells were fixed for 15 min at room temperature for 15 minutes using a fixation buffer (4% PFA, 0.02% sodium azide, and 0.1% Tween 20 in PBS) and washed with FACS buffer B (0.02% sodium azide and 0.1% Tween 20 in PBS). Subsequently, the cells were permeabilized at 4°C for 10 minutes in a permeabilization buffer (0.02% sodium azide, 0.1% Tween 20, and 10% dimethyl sulfoxide in PBS). The cells were washed and re-fixed in the fixation buffer for 5 minutes at RT. The staining and analysis procedures was the same as for surface protein detection. At least 30,000 cells were collected and 10% of cells with the highest GFP expression are shown.

### Tumor 3D spheroid formation and co-culture assay

Tumor cells were seeded in a Nunclon Sphera 96 well plate (ThermoFisher, 174925) at a density of 6,000 per well and cultured for 4-5 days to form tumor spheroid. Tumor 3D spheroid model was also established using HycopGel (COOP BIOTECH, CR01A0105) according to manufacturer’s instructions. The gel was incubated at 37°C for 10 minutes prior to use and mixed with the cell suspension at a 1:1 volume ratio to achieve a final cell concentration of 1×10L cells/ml. A 40 μl aliquot of the mixture was dispensed onto a pre-chilled 24-well culture plate and allowed to settle on ice for 5 minutes. Subsequently, 1 ml of 1×C Buffer, diluted in serum-free medium, was added to each well and maintained at 4°C for 15 minutes. The buffer was then replaced with complete culture medium, and cultured for 7 days. After tumor spheroid formation, ASO was transfected at a final concentration of 200 nM using JetPRIME. 20,000 macrophages were added after 48 hours. Fluorescence images were captured daily under the same parameter settings. The data are presented as the mean ± SEM of six independent experiments.

### Long-time microscopic photography

Long-time microscopic photography of co-culture system were captured on a high-speed confocal platform (Andor, Dragonfly 200). OVCAR3 cells were pre-transfected with 70 nM ASO 48 hours prior to experimentation. GFP labeled HEY and OVCAR3 cells were co-cultured with THP-1 induced macrophages staining with Cell Plasma Membrane Staining Kit with DiI (Beyotime, C1991). Time-lapse imaging was performed using Fusion software (v1.1.0, Andor) at an interval of 10 seconds per frame for a minimum duration of 120 minutes. The video was further processed using Imaris viewer (v9.0.1) and Adobe Premiere Pro (v22.0.0). Long-time monitoring splicing reporter cells and Live/Dead cells were conducted by Incucyte S3 Live-Cell Analysis System (Sartorius). ASO-treated reporter cancer cells were seeded in 6-well plate and allowed to adhere for 6 hours. Fluorescence images were captured for 48 hours with Incucyte S3. The fluorescence intensity was analyzed by Incucyte GUI (v2022B Rev1). We calculated the Green Object Count/Red Object Count ratio, normalized to t=0, and visualized the data using GraphPad Prism, presenting the mean ± standard deviation (SD). Three biological replicates performed. In the live and dead cells detection experiment, cancer cells were transfected with 70 nM ASO and macrophages were added 48 h later. The co-culture system was stained with Live and Dead Cell Double Staining Kit (Abbkine, KTA1001). The representative images at 8 h after co-culturation were shown.

### scRNA-seq of clinical samples and single cell alternative splicing analysis

Raw sequencing data of scRNA-seq was achieved from the GEO database. The raw reads were aligned to the human genome (hg38) by CellRanger (v7.1.0) with default parameter. Seurat (5.1.0) under R (4.4.1) was used for quality control, dimensionality reduction, and clustering. For each sample dataset, we filtered the expression matrix by the following criteria: 1) cells with total detected genes or UMI counts in either the top or bottom 2.5 % of the distribution were excluded; 2) cells with over 20 % of UMIs derived from the mitochondrial genome were excluded; 3) genes expressed in less than 3 cells were excluded. After filtering, 118,297 cells were used for the downstream analysis. The filtered object was normalized and then removed batch effects using SCTransform function. Principal component analysis (PCA) was performed on the scaled highly variable gene matrix, and the top 30 principal components (PCs) were selected for downstream clustering and dimensionality reduction. Cell clustering was performed using the FindClusters function (resolution = 0.8), yielding 26 distinct clusters. The resulting clusters were visualized using uniform manifold approximation and projection (UMAP). The differential expressed genes of each cell type were identified by the FindAllMarker function. Cell type was annotated by SingleR (v2.6.0) with manual correction. CellLcell communication analysis was conducted using the CellChat (v1.5.0). Pseudotime analysis was conducted using the monocle2 (v2.32.0).

For subcluster analysis of myeloid cells, the top 20 PCs were selected and resolution = 0.2 was used in FindClusters function. Each subgroup was annotated manually based on previous study^30^. M1 and M2 polarization scores for myeloid subgroups were computed by aggregating expression levels of previously established gene signatures ^31^. Pathway enrichment in each cell type was performed with the irGSEA package (v 3.3.2) using pre-defined gene sets in the MSigDB database. Pathway enrichment of macrophage subgroups in co-culture system was performed with the clusterProfiler package (v 4.12.4).

Alternative splicing analysis of the single cell was performed according to the SpliZ workflow^32^ using the parameter: SICILIAN=false, pin_S=0.1, pin_z=0.0, n_perms=100, libraryType=10X. The SpliZ or SpliZVD value of CD47 was extracted and merge with Seurat Object for visualization. All samples were divided into L dominant samples (SpliZVD value < 3) and S dominant samples (SpliZVD value ≥ 3). All cells were divided into Epi cells (CD47-L) (SpliZVD value < 3) and Epi cells (CD47-S) (SpliZVD value ≥ 3).

### scRNA-seq of co-culture system and scMicroAS-seq

OVCAR3 cells were transfected with 70 nM ASO-I10. After 48 h, 2×10^5^ OVCAR3 cell and 2 ×10^5^ THP-1 derived macrophage were co-cultured in 6 well plate (ThermoFisher, 174925) for 12 h. The three replicates in each group were mix together before captured by Perseus (10K Genomics). Libraries were prepared using Perseus Single Cell 3’ Reagent Kit (10K Genomics). Briefly, cells were co-encapsulated with 10K barcoded gel beads (containing UMIs and poly(dT) primers) using microfluidics. After reverse transcription and emulsion breaking, cDNA was amplified and processed through: (1) fragmentation/end repair, (2) SPRIselect size selection, (3) adaptor ligation, and (4) index PCR. Pooled libraries were sequenced on NovaSeq 6000 (PE150). The scRNA-seq data was processed by CellCosmo (10K Genomics) and further analyzed with Seurat similar to the procedures above. Cells with total detected genes or UMI counts less than 200 were excluded and genes expressed in less than 3 cells were excluded. The filtered object was normalized and removed batch effects using SCTransform function. The top 15 PCs were selected and resolution = 0.5 was used in FindClusters function. scMicroAS-seq was performed using the full-length cDNA before fragmentation. First-Round PCR Reaction used the targeted primer ∼50nt upstream the alternative splicing event and the anchor sequence in the P5 end. ChamQ SYBR Color qPCR Master Mix (Low ROX Premixed) (Q431-02, Vazyme) was used for amplification and monitor the Rn value reaches about 1.5 on QuantStudio 3 (Thermo Fisher Scientific). Dilute the product for 10 times and used for Second-round PCR with paired adaptors. After amplification for 8 cycles, the library was purified with AMPure XP beads. The library was sequenced on Illumina NovaSeq 6000 platform (PE150, 35M read-pairs/sample). The targeted amplification sequencing data was processed by CellCosmo (10K Genomics) and the mapped data were analyzed through the SpliZ workflow (v1.0dev). The SpliZ value of CD47 was extracted and merge with Seurat Object for visualization.

### Spatial Transcriptomics Library Preparation

Fresh tissues were frozen in OCT compound, and sections (RIN>7) were prepared using a Leica CM3050 cryostat. Brightfield imaging was performed on a Leica Aperio Versa 8 scanner (20×). Tissue permeabilization was optimized using the 10x Genomics Visium Tissue Optimization Kit (CG000238, 10x Genomics), testing variable durations to maximize mRNA capture while minimizing diffusion. The final optimized permeabilization condition was determined to be 6 minutes, confirmed by fluorescence signal intensity and spatial resolution. For library construction, sections (10 µm) were processed on Visium Gene Expression Slides following H&E staining, permeabilized for 6 minutes, and subjected to reverse transcription. Libraries were prepared per the 10x Genomics protocol (CG000239) and sequenced on NovaSeq PE150 (300M read-pairs/sample).

### Spatial transcriptome analysis

The samples used in the spatial transcriptome analysis were listed. Raw sequencing data of spatial transcriptome was demultiplexed and aligned to the human reference genome GRCh38-2020-A by SpaceRanger (v2.0.1). Count data was loaded into R (v4.4.1) using Load10X_Spatial function in Seurat (v5.1.0). Gene expression normalization, dimensionality reduction, clustering, and differential expression analysis were performed using Seurat. Spots with <100 detected genes were filtered out. Data were normalized via SCTransform, and 3,000 highly variable genes (HVGs) were selected for principal component analysis (PCA). For spot clustering, the first 30 PCs were used to build a graph, which was segmented with a resolution of 0.8. Differentially expressed genes (DEGs) were identified for each cluster using the FindAllMarkers function, with thresholds of fold change >2 and adjusted p-value<0.05. Cell type was annotated by SingleR with manual correction. The SpliZ value was calculated as above and added to the Seurat object. The distance and network between epithelial cells and macrophages were calculated and visualized using in house code, which was deposited in GitHub repository (https://github.com/PrinceWang2018/ST_Distance)

### Perturb-seq analysis

The public available Perturb-seq data (GSE264667) used a large-scale pooled CRISPRi screen targeting 2,285 essential genes with single-cell RNA sequencing experiments in Jurkat cells. The raw data was transformed into fastq format using parallel-fastq-dump (v0.6.7) with the parameter: –spilt-files. The reads were processed and mapped to human genome (hg38) by CellRanger (v7.1.0). The alternative splicing SpliZ value of CD47 in each cell was analyzed by SpliZ workflow as described above. The gRNA information of cells with corresponding barcodes was extracted from the h5ad file. The low-quality and outlier cells were filtered according to: 1) total count < 1000, 2) z_gemgroup_UMI <-1 or >1, 3) Z-score inside each target <-1.5 or >1.5. The medium SpliZ value of 134 core splicing factor were calculated ^33^. The splicing factors with over 10 cells detecting SpliZ value were further compared with non-target control and ranked. The p-value was calculated with unpaired two-tailed Student’s t-test. Moreover, the perturbations were visualized with minimum distortion embedding (MDE) using pyMDE (v0.1.18) according to previous method ^34^. Briefly, the expression profile of each sgRNA was z-normalized and the dataset was embedded using the SpectralEmbedding function of sklearn with n_components=20, affinity=’nearest_neighbors’, n_neighbors=7, eigen_solver=’arpack’. We then initialized preserve neighbors’ function of pyMDE with embedding_dim=20, n_neighbors=7, and repulsive_fraction=5.

### Quantitative proteomic profiling

Proteins were extracted from tissue samples using SDT lysis buffer (4% SDS, 100 mM DTT, 100 mM Tris-HCl, pH 8.0). The protein concentration was determined using a BCA Protein Assay Kit (P0012, BeyoTime). Protein digestion followed the FASP method ^35^. LC-MS/MS was performed using an Orbitrap Astral mass spectrometer coupled with a Vanquish Neo UHPLC system (Thermo Fisher Scientific) by Shanghai Bioprofle Technology Co., Ltd. Raw data was processed by DIA-NN (v1.9.0). We used the homo sapiens reference from UniProtKB (uniprotkb_taxonomy_id_9606_2024_08_27) to generate deep learning-based prediction of spectra with FASTA digest for library-free search / library generator and Deep learning-based spectra, RTs and lMs prediction function selected. Peptide length range was set to 7–30 and precursor charge range was set to 1–4, with the match-between-runs (MBR) function activated. Based on CD47-L specific sequence (KAVEEPLNAFKESKGMMNDE), we quantified the expression of CD47-L protein and CD47 total expression and calculated the ratio.

### Transcriptome Sequencing and Isoform Analysis

For short-read RNA-seq, total RNA was isolated from ASO treated OVCAR3 cells using TRIzol reagent (Invitrogen) according to the manufacturer’s protocol. RNA quality was assessed using an Agilent Technologies 2200 Bioanalyzer with the application of an RNA integrity number > 7.0. Poly(A) sequencing libraries were prepared using NEBNext Ultra RNA Library Prep Kit for Illumina (E7530L, NEB) following manufacturer’s recommendations. Adenylated mRNAs were isolated using oligo-d(T) magnetic beads (two rounds). The RNA-seq library was sequenced using Illumina NovaSeq 6000 (PE150) at Novogene. After obtaining paired-end reads and trimming adapter using TrimGalore (v0.6.1), clean reads were aligned to the hg38 genome with STAR (v2.2.0) and sorted with samtools (v1.9.0). Mapped reads were visualized with the Integrative Genomics Viewer (v2.16.0) and quantified using featureCounts in subread (v2.0.2). Differential expression was analyzed with DeSeq2 (v1.46.0). The cutoff was set as q < 0.05 and log fold change(logFC) > 1 or < -1. Gene and isoform expression level of TCGA-OV and GTEx ovary was achieved form UCSC Xena. The isoform switch events with consequence were analyzed using IsoformSwitchAnalyzeR (v2.2.0). PSI index used in pan-cancer analysis were achieved from TCGASpliceSeq database. 49 Immune checkpoint ligands were further analyzed.^36^ The raw sequencing data, gene expression FPKM and copy number variation of ovarian cancer cell lines were achieved from Cancer Cell Line Encyclopedia. PSI was calculated by Spliceseq (v2.1).

For long-read Iso-seq, the library was prepared according to the Isoform Sequencing protocol (Iso-Seq) using the Clontech SMARTer PCR cDNA Synthesis Kit and the BluePippin Size Selection System protocol as described by Pacific Biosciences (PN 100-092-800-03). The library was sequenced on PacBio Sequel II (∼23M reads/sample). Sequence data were processed using the IsoSeq (PacBio, v3.8.0) software. Circular consensus sequence (CCS) was generated from subread BAM files, parameters: --min-rq 0.9 –min-passes 1 -j 22. Barcode and primer sequences were identified using lima function with parameters: --isoseq –peek-guess. Poly(A) and concatemer were removed using refine function. The subreads were mapped to the hg38 reference using pbmm2 (v1.9.0) and convert to fastq format using bam2fasta function in pbtk (v3.5.0). Isoform quantification was performed using salmon and analyzed with IsoformSwitchAnalyzeR (v2.2.0). Alternative events were analyzed with SUPPA (v2.3.0).

### MicroAS-seq

We developed a targeted multiplex amplification library construction protocol to detect the micro-exons skipping. Briefly, targeted Round1 PCR primers with dual barcode and variable spacers were used to amplify the alternative splicing region of CD47. Variable-length spacers were employed to mitigate nucleotide composition bias and enhance sequencing quality. A 10-fold diluted cDNA aliquot served as the template for PCR amplification. Quantitative PCR amplification was performed using ChamQ SYBR Color qPCR Master Mix (Low ROX Premixed; Q431-02, Vazyme) on a QuantStudio 3 system (Thermo Fisher Scientific), with amplification terminated when the normalized reporter signal (Rn) reached ∼1.5. The primary PCR product was diluted 10-fold and subjected to a second-round PCR (8 cycles) with adapter-specific primers. Libraries were purified using AMPure XP beads (Beckman Coulter) and sequenced on an Illumina NovaSeq 6000 platform (150 bp paired-end, 5 million read-pairs per sample). The custom pipeline has been deposited in the Github repository (https://github.com/PrinceWang2018/MicroAS-seq).

### LACE-seq2

To improve library compatibility and tailor the protocol for cancer cell applications, we optimized the original LACE-seq method^37^, renaming the updated version LACE-seq2. LACE-seq2 identify RNA binding protein targets using UV crosslinking and linear amplification of complementary DNA ends method. Briefly, 5 μg HNRNPA1 and IgG antibody were pre-incubated with 50 μl pre-washed Protein A/G magnetic beads (Thermo Scientific, AM9624) at 4 □ overnight. 500,000 HEY cells were crosslinked with UV-C in CL-1000 Ultraviolet Crosslink for 350 mJ twice in 6 cm dish on ice. Collect and lysis the cell on ice for 10 min. Add 10 μl SUPERase•In RNase Inhibitor and remove of genomic DNA with 40 μl RQ1 RNase-Free DNase at 37 □ for 3 min at 1200 rpm. Incubate the antibody binding protein A/G beads with the cell lysis supernatant. Rotate the mixture for 2 h at 4 L. 0.3 U Micrococcal Nuclease (Thermo Scientific, EN0181) were used for RNA fragmentation and incubated at 37 □ for 12 min. Stop the reaction with 20 mM EGTA. The dephosphorylation RNA fragments were ligated to biotin-labeled 3’linker using T4 RNA Ligase 2, truncated (M0242S). Both gentle washing off and on the magnetic stand were utilized. After reverse transcription with Superscript II Reverse Transcriptase (Thermo Scientific, 18064014), the cDNA was treated with Exonuclease I (NEB, M0293S) at 37 □ for 1h and RNase H (NEB, M0297S) at 37 □ for 30 min. Poly(A) tailing was performed with terminal transferase (NEB, M0315S) at 37 □ for 10 min. The dual-index (P7;P5) adapters are used for amplification and construction of the sequencing library. The fragments with lengths between 230 bp and 600 bp were selected for sequencing. Sequencing was performed with Illumina NovaSeq X plus platform using PE150 strategy (Novogene).

### Crosslink sites identification and motif analysis

The adapters of raw sequencing data were removed using Cutadapt with the parameter: -j 8 -q 30,0 --trim-n --max-n 0.25. The poly(A) and poly(T) sequences were removed with: -m 18 -n 2 -j 8. The quality of the reads was determined with fastqc. rRNA was removed with bowtie2 through aligning to human rRNA reference. To mitigate the impact of poly(A) on sequencing quality, only the first read in paired reads was aligned to the human genome (hg38) using STAR (v2.2.0) with default parameter. The aligned reads were indexed using samtools (v1.9.0) and converted bed format using bamToBed function in bedtools (v2.31.0). Two technical replicates were merged and call peaks using Piranha (v1.2.1) with the parameter: -s -p 0.001 -b 20 -d ZeroTruncatedNegativeBinomial -v. The peaks in the IgG group were removed and the remaining peaks were annotated using ChIPseeker (v1.40.0). For motif analysis, peaks were first extended 30nt on both sides. The de novo binding motif was identified using HOMER (v4.11) with the parameter: -len 6 -S 10 -rna -p 4. The motif location was detected using scanMotifGenomeWide.pl and visualized using Deeptools (v3.1.3). The CLIP-seq and eCLIP-seq data were aligned according to previous studies, and peaks were called using Piranha for comparison. The library variety was evaluated with preseq (v3.2.0). The custom pipeline used to analyze the LACE-seq2 data has been deposited in the Github repository (https://github.com/PrinceWang2018/LACE-seq2).

## Notes

### Competing Interest Statement

The authors have declared no competing interest.

