## Supplemental Figure 1 and 2 for "Isoform switch of CD47 provokes macrophage-mediated pyroptosis in ovarian cancer"

### Supplementary Figures

#### Supplementary Figure 1, related to Figure 1

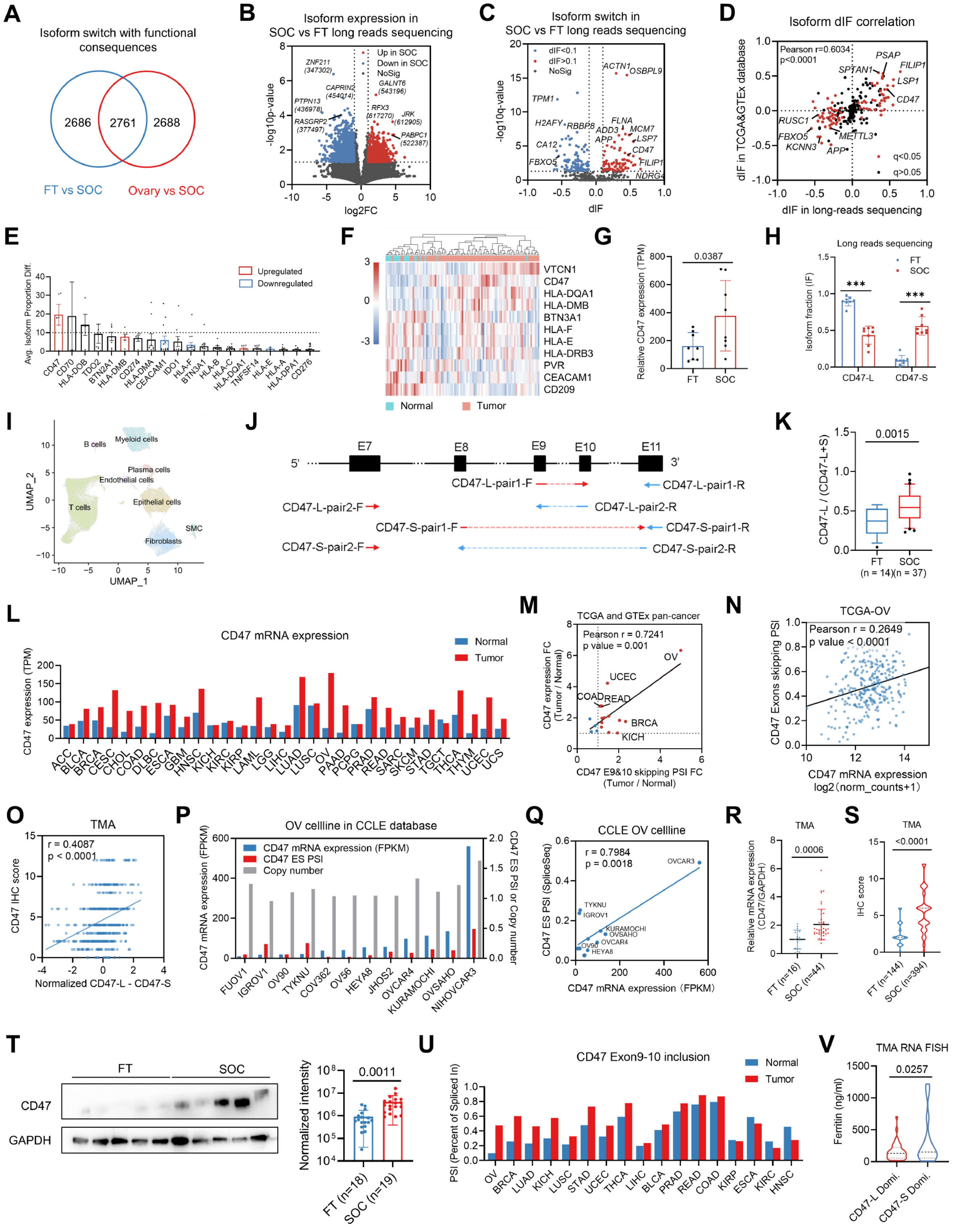

**Supplementary Figure 1. Pan-cancer analysis of immune checkpoint CD47 and its isoforms. (A)** The Venn diagram demonstrates the common targets analyzed in ovarian cancer with consequence of isoform switching, using normal fallopian tube umbrella tissue or ovarian tissue as control. **(B)** Volcano plot of differential expressed isoforms in long reads sequencing data of 8 pairs of SOCs and FTs. **(C)** Volcano plot of differential Isoform fraction (dIF) of the Isoform switch events. **(D)** Correlation analysis of the isoform switch events in TCGA-OV and GTEx database and our long reads sequencing data. **(E)** Average value of differential isoforms proportion of each immune checkpoint. Data was shown as mean with SEM. **(F)** Heatmap of protein expression pattern of immune checkpoints with significant changes in SOCs and normal tissues from CPTAC database. **(G, H)** CD47 mRNA expression **(G)** and isoform fraction **(H)** of FTs and SOCs in long reads sequencing data. **(I)** UMAP projection of all cells from scRNA-seq data of 5 FTs and 5 SOCs, colored by cell types. **(J)** Schematic diagram of the isoform-specific qPCR primer design scheme for detecting CD47-L and CD47-S. **(K)** qPCR verification of CD47-L ratio in FTs (n = 14) and SOCs (n = 37). **(L)** Pan-cancer analysis of CD47 mRNA expression in TCGA-OV comparing with normal tissue in GTEx database. **(M)** Differential fold change of CD47 mRNA expression and micro-exon skipping PSI comparing with normal samples in TCGA and GTEx database. **(N)** The correlation analysis of CD47 exon 9 and 10 skipping PSI and mRNA expression in ovarian cancer samples in TCGA-OV. **(O)** Correlation between IHC score of CD47 protein and difference between CD47-L and CD47-S immunofluorescence intensity. **(P)** CD47 mRNA expression, CD47 multi-exon skipping PSI index, and CD47 copy number in CCLE ovarian cancer cell lines. mRNA expression was normalized with FPKM. **(Q)** The correlation analysis of CD47 exon 9 and 10 skipping PSI and mRNA expression in ovarian cancer cell lines in CCLE database. **(R and S)** Relative CD47 mRNA expression **(R)** and IHC score **(S)** of FTs and SOCs in TMA array. **(T)** Representative western blot image and quantification of CD47 total protein expression in FTs (n = 18) and SOCs (n = 19). **(U)** Pan-cancer analysis of CD47 exon 9 and 10 skipping level (PSI) in TCGA-OV comparing with normal tissue in GTEx database. **(V)** The relation of CD47 isoforms expression pattern with ferritin concentration based on the RNA FISH quantification result of Qilu cohort.

**Supplementary Figure 2, related to Figure 2**

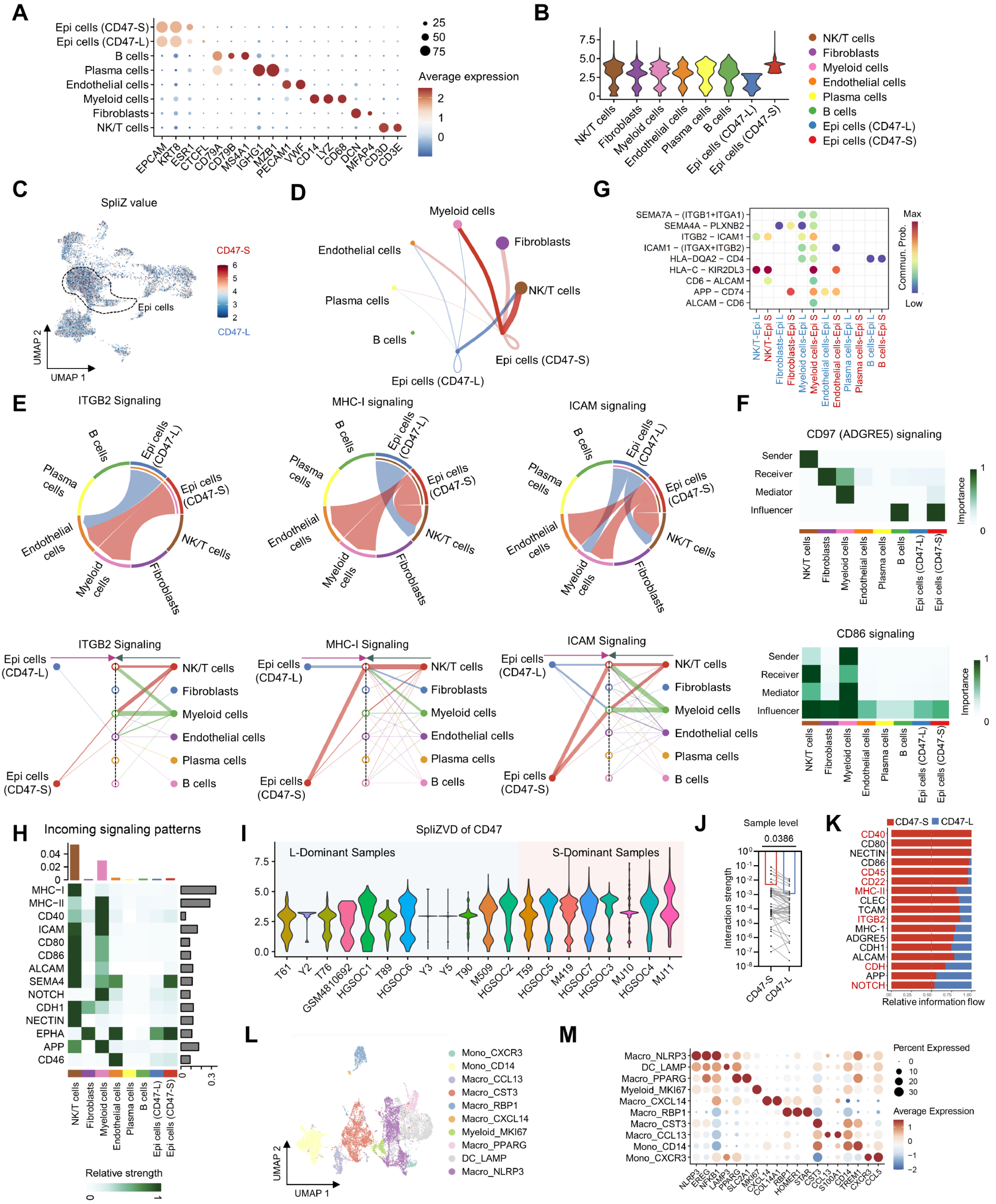

**Supplementary Figure 2. Cell commination analysis of CD47-L and CD47-S epithelial cells or CD47-L/S dominant samples. (A)** Normalized expression of marker genes in each cell cluster. **(B)** Violin plot of the SpliZVD value in CD47-L and CD47-S epithelial cells, myeloid cells, NKT cells, B cells, pre-B cells and endothelial cells. **(C)** UMAP projection of SpliZ value of all cells. Dashed circles indicate epithelial cell subgroup. Red, CD47-S cells; Blue, CD47-L cells. **(D)** Cell-cell communication network illustrating interactions between myeloid cells, fibroblasts, endothelial cells, NKT cells, B cells, pre-B cells, CD47-L and CD47-S epithelial cells. The thickness of the connections represents the strength of the interactions. **(E)** Circos plot and network diagram indicated the interaction ratio of ITGB2, MHC-1, ICAM signaling among CD47-L and CD47-S epithelial cells, myeloid cells, NKT cells, and endothelial cells. **(F)** CD97 (ADGRE5) and CD86 signaling heatmap depicting the roles of sender, receiver, mediator, and influencer cells. The color gradient reflects the interaction importance. **(G)** Dot plot showing the communication probability (Commun. Prob.) of ligand-receptor interactions between CD47-L/S epithelial cells and others. The color gradient represents communication strength, ranging from low (blue) to high (red), and dot size indicates interaction significance. **(H)** Heatmap of incoming signaling patterns showing relative signaling strength (color gradient) for various pathways across cell types. The bar plot on the right quantifies the overall relative signaling strength for each cell type. **(I)** SpliZVD value of CD47 in scRNA-seq data from 20 tissues. Average SpliZVD value over 3 was defined as S-dominant samples (red) while others were L-dominant samples (blue). **(J)** Quantification of the interaction strength of CD47-L/S dominant samples with other cell type. **(K)** Bar plot illustrating the relative information flow of ligand-receptor interactions mediated by CD47-S (red) and CD47-L (blue) isoforms. **(L)** UMAP projection of myeloid subgroups. **(M)** Dot blot of normalized expression of marker genes in each myeloid subgroup.
